# Gut bacterial metabolite imidazole propionate potentiates Alzheimer’s disease pathology

**DOI:** 10.1101/2025.06.08.657719

**Authors:** Vaibhav Vemuganti, Jea Woo Kang, Qijun Zhang, Eric R. McGregor, James R. Hilser, Ruben Aquino-Martinez, Sandra Harding, Joseph Lawrence Harpt, Katharina R. Beck, Hailey Bussan, Jessamine F. Kuehn, Yuetiva Deming, Sterling C. Johnson, Sanjay Asthana, Henrik Zetterberg, Kaj Blennow, Corinne D. Engelman, Hooman Allayee, Rozalyn M. Anderson, Tyler K. Ulland, Fredrik Bäckhed, Barbara B. Bendlin, Federico E. Rey

## Abstract

The gut microbiome modulates metabolic and neurovascular processes implicated in Alzheimer’s disease and related dementias (ADRD), but the underlying mechanisms remain unclear. Here, we identify the bacterial metabolite imidazole propionate (ImP) as a modifier of ADRD pathology. In a cohort of 1,196 cognitively unimpaired adults, higher plasma ImP levels were associated with lower preclinical cognitive scores and biomarkers of ADRD, both cross-sectionally and longitudinally. Fecal metagenomic analysis linked putative ImP producers to ADRD phenotypes. Genome-wide integrative analysis revealed a locus on chromosome 12 associated with both plasma ImP levels and AD risk in humans, supporting a host genetic contribution to ImP regulation and a causal role of this metabolite in AD. In mice, chronic ImP administration exacerbated AD-like pathology. Mechanistically, ImP impaired brain endothelial barrier and promoted tau hyperphosphorylation in primary neurons, an effect blocked by glycogen synthase kinase-3β inhibition. Together, our study links ImP to hallmarks of neurodegeneration and suggest that targeting ImP may represent a potential strategy to modify ADRD risk.

## MAIN

Alzheimer’s disease (AD) is a progressive neurodegenerative disorder characterized by amyloid-β accumulation, tau pathology, and neuronal loss^1^. Aging and the apolipoprotein E (*APOE*) ε4 allele^2^ remain the strongest risk factors for AD, but additional variables, including sex and vascular risk factors modulate disease severity^2^. More than 55 million people are living with dementia worldwide, with AD accounting for 60–70% of cases. Current treatment approaches primarily focus on slowing disease progression and alleviating symptoms. While current anti-amyloid therapy approaches offer promise, identifying early biomarkers and modifiable risk factors remains a priority for reducing dementia burden.

Multiple lines of evidence suggest that alterations in the gut microbiota may precede or co-occur with AD pathology^3–5^. Gut microbes produce a wide array of metabolites that modulate host physiology and may influence neurodegenerative processes^4^. Many gut microbial metabolites enter the bloodstream, where they can influence central nervous system (CNS) function indirectly via peripheral signaling or directly by crossing the blood-brain barrier (BBB). Previous studies have identified several bacterial metabolites associated with AD^4,6^, implicating gut bacterial metabolism as a potential contributor to neurodegeneration.

Imidazole propionate (ImP) is a metabolite produced during anaerobic metabolism of histidine by bacteria expressing the enzyme urocanate reductase (UrdA)^7,8^, which catalyzes the reduction of urocanate during anaerobic respiration. Elevated blood levels of ImP have been linked with multiple dementia risk factors, including type II diabetes (T2D)^9,10^, hypertension^11,12^, atherosclerosis^13^, and chronic kidney disease^14,15^, suggesting potential detrimental effects on host health. However, the impact of ImP on brain function and AD pathophysiology remains largely unexplored.

Building on evidence linking ImP to cardiometabolic conditions, including insulin resistance, endothelial impairment, and vascular inflammation, all of which elevate risk for ADRD, we hypothesized that ImP directly perturbs neurovascular and neuronal pathways relevant to cognitive aging. Specifically, given its known capacity to alter host kinase signaling and endothelial function, we posited that sustained ImP exposure compromises blood-brain barrier integrity and promotes tau-associated neurodegenerative processes. To address this, we integrated cognitive assessments^16^, neuroimaging^17^, fluid and imaging biomarker, and multiomic data^18^ from the Wisconsin Alzheimer’s Disease Research Center (ADRC), Wisconsin Registry for Alzheimer’s Prevention (WRAP)^19^ and Microbiome in Alzheimer’s Risk Study (MARS)^3,20^. We examined relationships between circulating ImP, the abundance of bacteria encoding UrdA homologues, cognitive performance, and AD pathology. To test the causal role of ImP in AD, we combined a genetics approach in humans with transgenic mouse models that capture key features of AD pathology and *in vitro* cell culture models. Together, our findings suggest that ImP disrupts neurovascular integrity and exacerbates AD-related neuropathology and provide mechanistic insights into how this bacterial metabolite contributes to cognitive aging and neurodegeneration.

## RESULTS

### Imidazole propionate is associated with lower preclinical cognitive scores

Plasma levels of ImP measured in 1,196 cognitively unimpaired individuals from the Wisconsin ADRC and WRAP cohort^19^ (mean age of 61.2 years), were tested in relation to ADRD-related outcomes which included cognitive scores, cerebrospinal fluid (CSF) and plasma biomarkers of ADRD pathology. **Extended Data Table.1** and **Extended Data Figure.1** summarize key demographic and clinical characteristics of the study cohort. Initial assessments revealed a positive relationship between age and plasma ImP levels (**Fig. 1a**). We also found plasma ImP levels were elevated in males compared to females (**Fig. 1b**). Thus, we used the ordinary least squares (OLS) multiple linear regression approach to examine the relationship between ImP and cognitive function, controlling for age, sex, BMI, *APOE* ε4 carrier status, and the age gap between plasma sample collection and cognitive assessment. We found that higher ImP was significantly associated with lower cognitive performance (**Fig. 1c and Extended Data Figure. 2**) as indexed by modified Preclinical Alzheimer’s Cognitive Composite (mPACC3) scores. The mPACC3 was adapted to address sampling limitations during the COVID-19 pandemic with remote assessments^21^. These results suggest that ImP is potentially an early, pre-clinical, independent contributor to cognitive function. Since ImP was previously associated with comorbidit ies of ADRD such as T2D, hypertension, atherosclerosis, and chronic kidney disease, we examined the extent to which these conditions contribute to observed ADRD measurements described above. We used the LIfestyle for BRAin health index (LIBRA)^22,23^, which includes risk factors such as midlife obesity, smoking, coronary heart disease, physical inactivity, chronic kidney disease, diabetes, midlife hypercholesterolemia, hypertension, and depression. We observed a significantly positive association between LIBRA scores and plasma ImP levels (**Fig. 1d**). Thus, we performed mediation analysis for the effects between ImP and mPACC3 cognitive scores, identifying that only 13-16% of the indirect effect to be mediated by ImP between LIBRA and mPACC3 results (**Extended Data Table 2**). Next, we employed the LIBRA index (**Fig. 1d**) as an additional covariate when testing the relationship between ImP and mPACC3. This showed that while lifestyle and ADRD risk factors explain a portion of the association, ImP remained the primary contributor to the observed effect (**Extended Data Figure. 2c**), suggesting that ImP acts largely as an independent factor rather than exerting its influence indirectly through comorbidities. These results suggest that ImP is associated with pre-clinical cognitive scores (**Fig. 1 and Extended Data Figure. 2**). There was no significant relationship between ImP and [^18^F]-fluorodeoxyglucose (FDG) standard uptake value ratio (SUVR) of the frontal, temporal, parietal, and occipital lobes (*N* = 168) (**Extended data Figure. 3a**). However, when stratified by sex (**Extended Data Figure. 3b & c**), a significant negative association was detected in males for the frontal lobe (β = −0.069, *P* = 0.043, *N* = 51) and parietal lobe (β = −0.083, *P* = 0.037, *N* = 51), while no significant relationship was observed in the temporal lobe (β = −0.04, *P* = 0.16, *N =* 51) and the occipital lobe (β = −0.04, *P* = 0.26, *N =* 51) after controlling for covariates (**Extended data Figure 3c**). The effect observed in frontal lobes of males was not statistically significant when high ImP outlier was not accounted in the regression (**Extended Data figure. 3d**).

**Figure 1.**
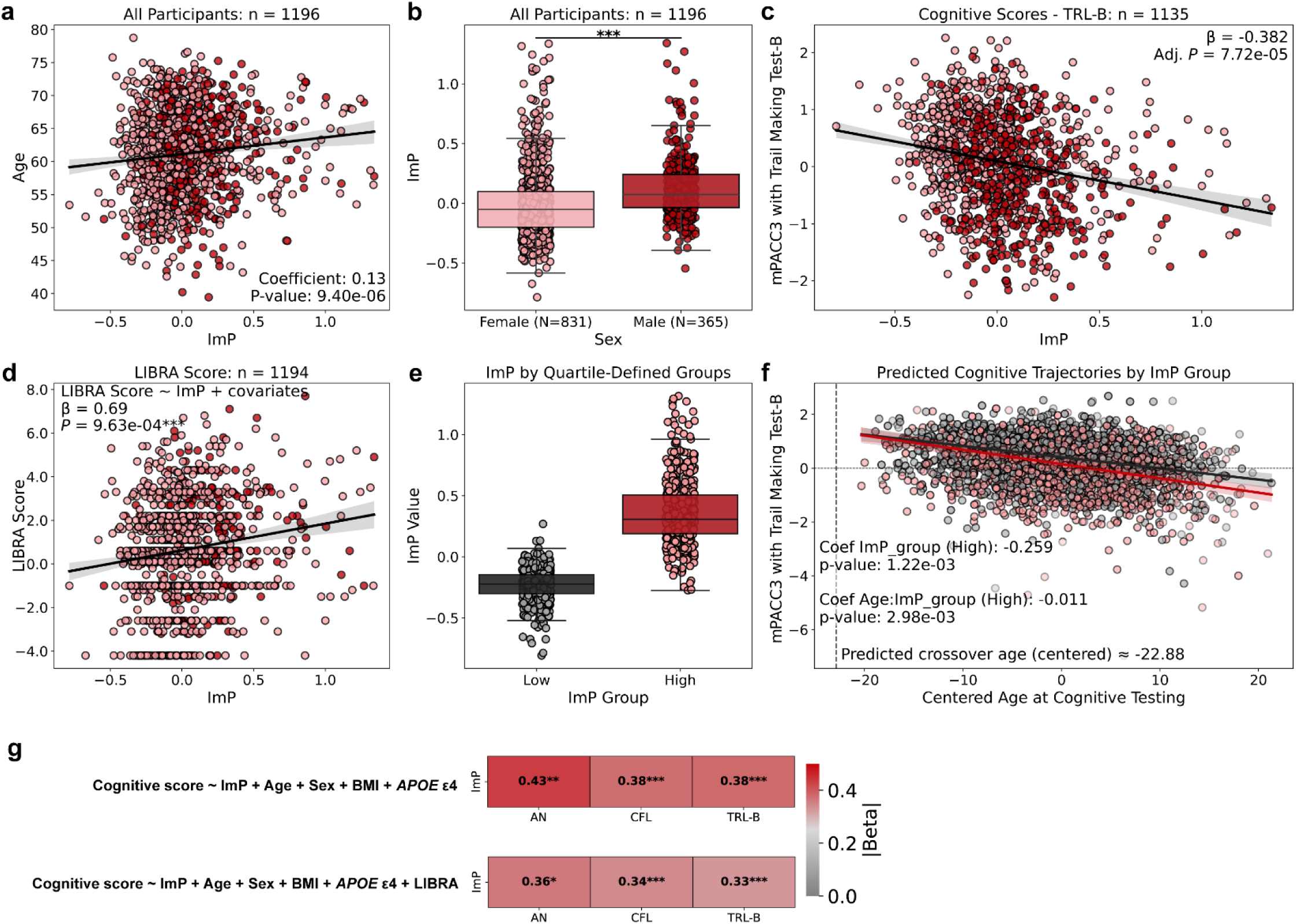
Plasma ImP is associated with preclinical cognitive trajectories. a, Plasma ImP levels increase with age. **b,** Plasma ImP levels are higher in males than females (*P* < 0.001, Mann–Whitney *U* test). **c,** Higher plasma ImP levels are associated with lower mPACC3 scores cross-sectionally. **d,** Plasma ImP levels are positively associated with LIBRA scores. **e,** Distribution of normalized ImP levels across the highest ImP (red) and lowest ImP quartiles (gray). **f,** Modeled cognitive trajectories over age for mPACC3 scores, with the vertical dashed line indicates predicted crossover age and the horizontal dashed line denotes zero on the scaled y-axis, comparing top (red) and bottom (gray) ImP quartiles. **g,** Summary heat-maps showing absolute β coefficients for outcomes attributable to ImP, estimated using ordinary least-squares (OLS) regression models with and without adjustment for LIBRA.

**Figure 2.**
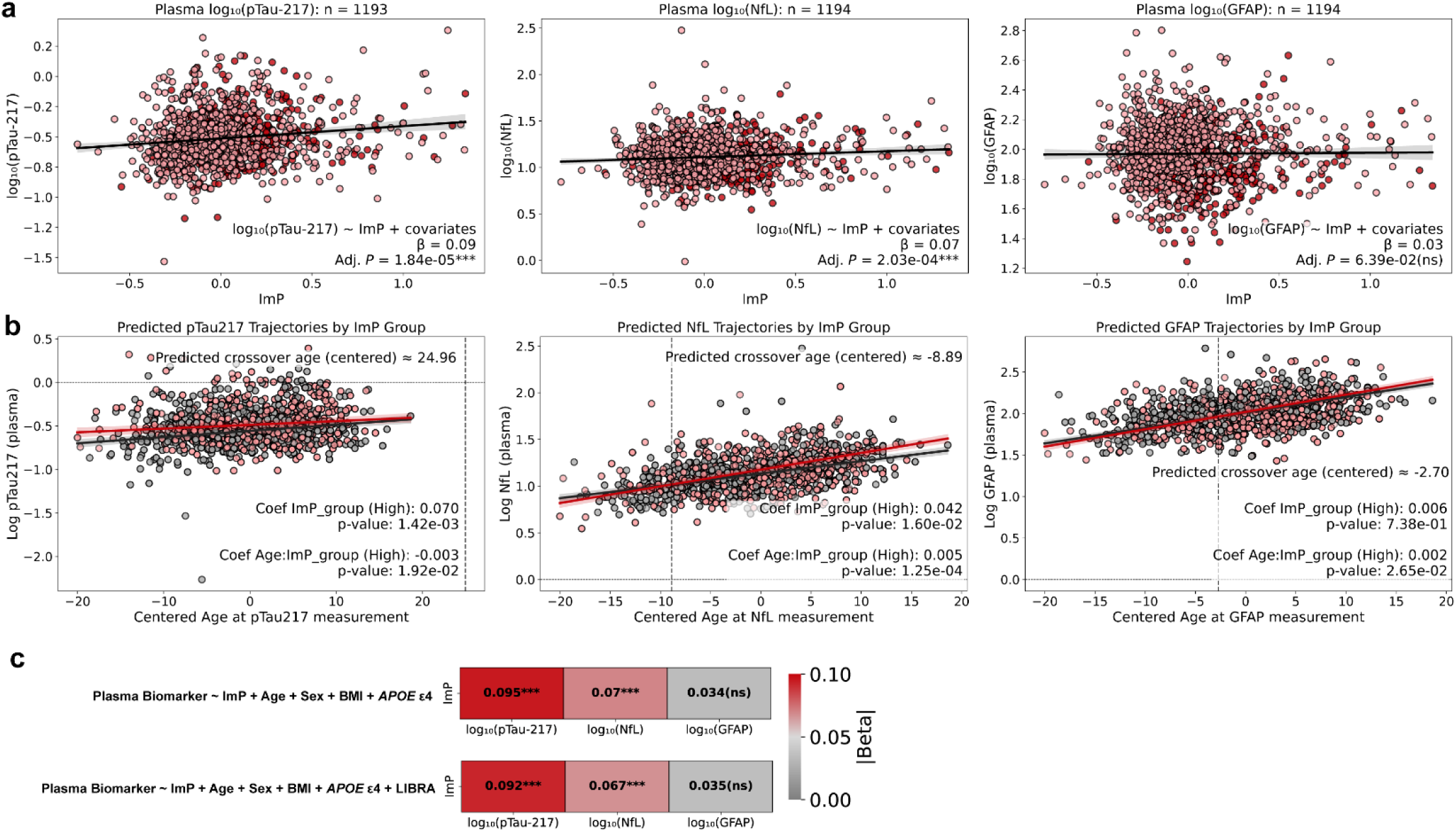
Plasma ImP is associated with ADRD plasma biomarkers. a, Cross-sectional associations between plasma ImP levels and ADRD biomarkers, pTau-217, NfL and GFAP. **b,** Modeled age-related trajectories for plasma pTau-217, NfL, and GFAP, with the vertical dotted line indicating predicted crossover age and the horizontal dotted line marking zero on the scaled y-axis, comparing highest and lowest ImP quartiles. **c,** Summary heat maps displaying β coefficients from OLS regression models relating ImP to plasma biomarkers, estimated with and without LIBRA adjustment.

**Figure 3.**
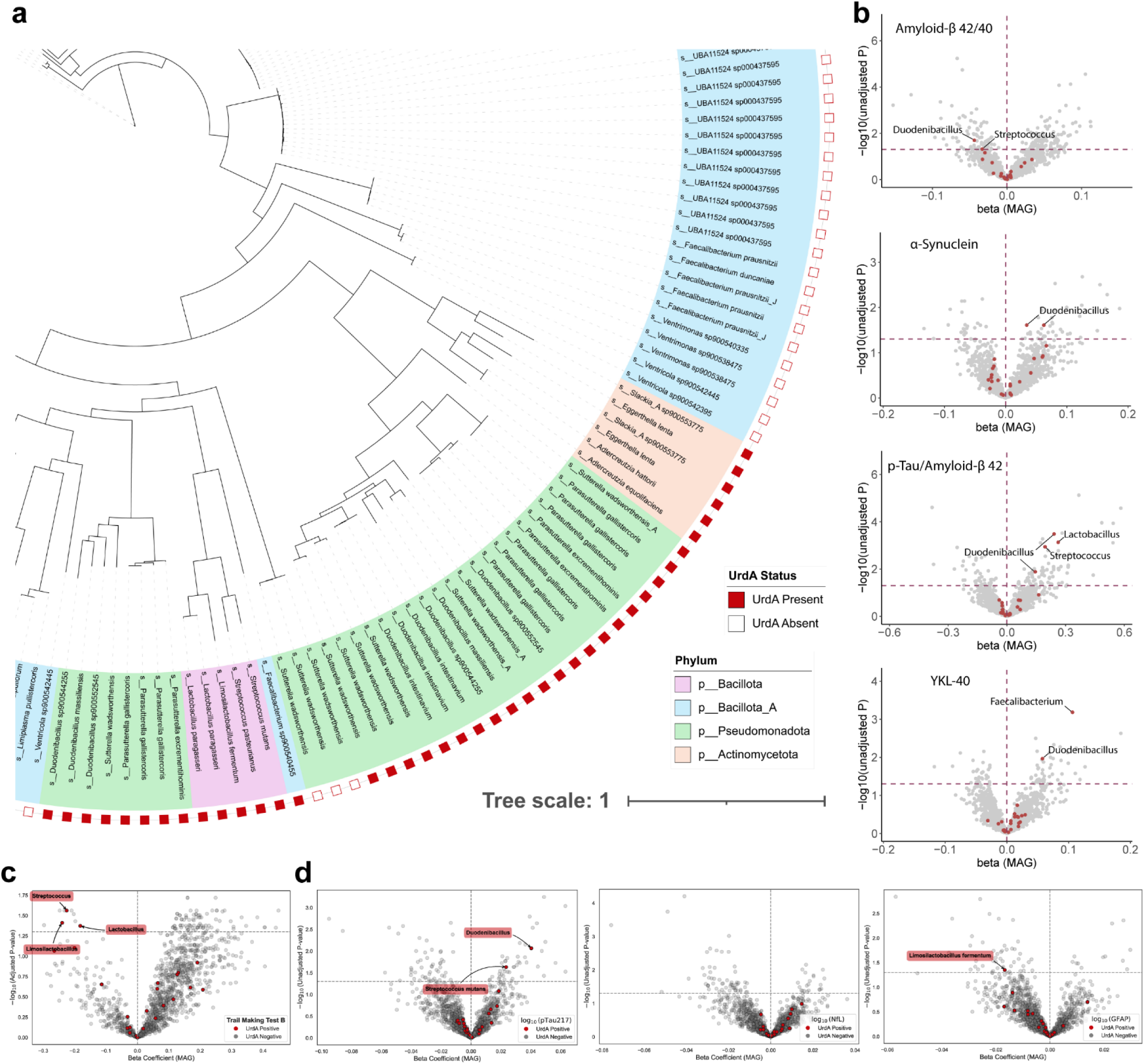
Gut bacteria encoding putative *urdA* orthologs are associated with cognitive performance and ADRD-related biomarkers. a, Section of a phylogenetic tree depicting the various MAGs within MARS cohort (N = 294). Each terminal branch represents a species, colored by phylum. The outer layer of the tree indicates the presence (red) or absence (white) of the *urdA* gene. **b,** Volcano plots displaying associations between abundance of *urdA*+ MAGs and CSF biomarkers of ADRD. **c,** Volcano plots depicting associations between abundances of *urdA*+ MAGs and cognitive performance (mPACC3) scores. **d,** Volcano plots representing associations between abundances of *urdA*+ MAGs and plasma biomarkers of ADRD.

We evaluated the relationship between ImP and longitudinal cognitive decline by using longitudinal mPACC3 data from multiple visits (**Extended Data Table. 3** & **Extended Data Figure. 4**). We stratified participants into quartiles based on the pooled distribution of plasma ImP levels across all participants and timepoints. Analyses focused on individuals in the highest and lowest ImP groups (**Extended Data Figure. 4**, **Fig. 1e**). Although the number of observations per participant varied, all available timepoints were included in the mixed-effects models, which accommodate unbalanced longitudinal data. We used mixed-effects models to examine differences in both baseline levels and age-related changes in cognitive performance. To standardize the time scale across participants, we centered age at the sample level, enabling the assessment of age-related trajectories while accounting for varying visit times. This approach allowed us to test whether cognitive decline differed by ImP group while controlling for relevant covariates and within-subject correlations. Participants in the highest ImP quartile exhibited significantly lower baseline cognitive scores and steeper age-related decline compared to those in the lowest quartile, across all three mPACC3 (**Fig. 1f and Extended Data Figure. 2b**). For example, compared with the low-ImP group, the high-ImP group showed a lower baseline score on mPACC animal naming (β = −0.347, *P* = 0.005) and exhibited a steeper negative age-related trajectory (interaction β = −0.022, *P* < 0.001). Similar patterns were observed for mPACC letter fluency (β = −0.252, *P* = 0.005; interaction β = −0.017, *P* < 0.001) (**Extended Data Figure. 2b**) and mPACC trail making test-B (β = −0.252, *P* = 0.002; interaction β = −0.014, *P* < 0.001) (**Fig. 1f**). The divergence in cognitive trajectories between the high- and low-ImP groups widened over time. Model projections indicated crossover points, where the high group’s cognitive performance declined below that of the low-ImP group, at approximately 15.5 years (animal naming), 14.6 years (letter fluency), and 18.2 years (trail making test-B), before the samples’ mean age. These findings suggest that higher ImP levels are associated not only with worse baseline performance but also with earlier and more rapid cognitive decline.

**Figure 4.**
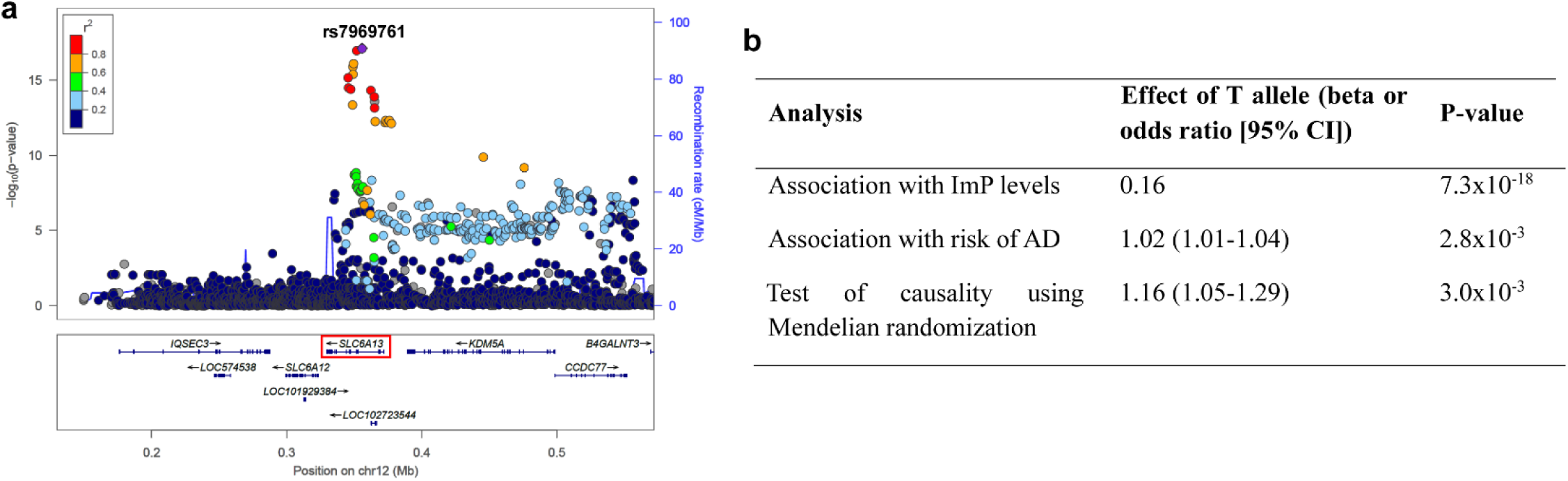
Genetic evidence supporting causal association between elevated plasma ImP levels and AD risk. a, The lead variant, rs7969761, maps to an intronic region of the *SLC6A13* gene (red box), with the T allele being associated with increased ImP levels. **b,** Effect sizes are shown with respect to the T allele of rs7969761. Causal association of 1 standard-deviation increase in normalized ImP levels with risk of AD was evaluated using a Wald ratio test with published effect sizes for rs7969761.

### ImP is associated with biomarkers of ADRD

The observed associations suggest a shift toward neurodegenerative pathology in individuals with elevated ImP levels. Plasma ImP showed a significant positive relationship between plasma ImP and NfL (neurofilament light chain) in cerebrospinal fluid (CSF) (Spearman, R = 0.2; Adj. *P* = 0.003, *N* = 224) a well-established marker of neurodegeneration^24^ (**Extended Data Fig. 5**). Consistent with this, plasma ImP was also positively associated with plasma NfL (β = 0.07, *P* = 2.03e-04) (**Fig. 2a**). Moreover, plasma ImP correlated with plasma pTau-217 (β = 0.09, *P* = 1.84e-05) (**Fig. 2a**) a biomarker of AD pathology^25^. A trend was also observed between ImP and plasma levels of glial fibrillary activation protein (GFAP), a biomarker of astrocyte activation^26^ (β = 0.03, *P* = 6.39e-02) (**Fig. 2a**), further suggesting that elevated ImP associates with astroglial activation characteristic of neurodegenerative disease.

We examined the longitudinal association between ImP status and pTau-217. Individuals in the high-ImP group showed significantly higher baseline levels of pTau-217 (**Fig. 2b**, β = 0.070, *P* = 0.001), indicating an increased Alzheimer’s-related biomarker burden. Although pTau-217 levels increased with age overall (positive slopes), the high-ImP group showed a slightly attenuated rate of age-related increase (interaction β = −0.003, *P* = 0.019). Of note, the predicted crossover in pTau-217 between the high- and low- ImP groups occurred around +25.0 years on the centered age scale, suggesting that individuals with high ImP reach elevated biomarker levels earlier, and may experience a plateau phase as they age (**Fig. 2b**). This pattern supports the hypothesis that high ImP may reflect an earlier pathological shift in both cognitive and biomarker trajectories.

We also examined the longitudinal relationship between ImP status and NfL, using mixed-effects modeling. Participants in the high-ImP group showed significantly elevated baseline levels of log-transformed NfL compared to the low-ImP group (β = 0.042, *P* = 0.016), consistent with greater neuronal injury burden. A significant age-by-ImP interaction (β = 0.005, *P* < 0.001), a steeper age-related increase in NfL among individuals with high ImP compared to those in the low-ImP group. The modeled crossover point, where NfL levels in the high-ImP group exceeded those in low-ImP group, occurred at a centered age approximately −8.9 years, supporting an earlier and accelerated trajectory of neurodegeneration (**Fig. 2b**).

At baseline, plasma GFAP levels did not differ by ImP status (β = 0.006, *P* = 0.74). However, the high-ImP group showed a significantly steeper increase in GFAP with aging (β = 0.002, *P* = 0.03), with divergence between groups emerging around a centered age of −2.70 years, consistent with earlier and accelerated AD-related astrocyte reactivity (**Fig. 2b**). This pattern mirrors findings for cognitive decline, pTau217, NfL, and GFAP (**Fig. 1g** & **2c**), agnostic to lifestyle factors (**Extended Data Figure. 6**), suggesting that elevated ImP may indicate early and progressive neurobiological vulnerability across multiple Alzheimer’s-related markers.

### Metagenome-assembled genomes (MAGs) encoding putative urocanate reductase are associated with Alzheimer’s disease

We assessed the gut microbiome’s capacity to generate ImP by quantifying the relative abundance of bacterial species harboring putative orthologs of the enzyme urocanate reductase (UrdA) and examined its association with AD biomarkers. We generated metagenome datasets from fecal samples (n=294) collected from individuals enrolled in the MARS cohort (**Extended Data Table 4**). Sequencing data from each sample were individually assembled, and contigs were clustered into metagenome-assembled genomes (MAGs), resulting in a total of 1,464 high-quality MAGs (>90% completeness and <5% contamination). We detected the presence of putative *urdA* orthologs in several MAGs including *Turicimonas muris, Eggerthella lenta, Faecalibacterium prausnitzii, Streptococcus mutans, Streptococcus pasteurianus, Duodenibacillus massiliensis, Lactobacillus paragasseri, Adlercreutzia hattorii, Sutterella wadsworthensis, Sutterella megalosphaeroides, Limosilactobacillus fermentum* and *Adlercreutzia equolifaciens* (**Fig. 3a**, **Extended Data Fig. 7a**), all of which were confirmed to have the tyrosine (Y) or methionine (M) residue at 373 in the FAD binding site, which are key for filtering enzyme specificity towards urocanate by excluding false positive hits for fumarate reductases^7,8,10^ (**Extended Data Fig. 7b)**. Some of these taxa have previously been identified as ImP producers^10,12,27^. We examined associations between abundances of fecal *urdA^+^* MAGs and plasma ImP levels and found only one bacterial taxa (*Streptococcus pasteurianus*) significantly positively associated with ImP (**Extended Data Figure. 7c**). We then applied multiple linear regression models to identify MAGs whose abundances were significantly associated with CSF biomarkers related to AD. MAGs positive for *urdA* were significantly associated with AD biomarkers, namely amyloid-β 42/40 ratio, α-synuclein, p-Tau/ amyloid-β 42 ratio, and YKL-40 (**Fig. 3b**). Furthermore, *urdA^+^* MAGs from the genera *Limosilactobacillus, Lactobacillus* and *Streptococcus* were negatively associated with cognitive scores spread across the mPACC3 tests (**Fig. 3c**). These results suggest that the abundance of gut bacterial taxa potentially harboring a key gene for ImP production is associated with AD. However, only *Duodenibacillus* and *Streptococcus mutans* were associated with pTau-217 (**Fig. 3d**), no *urdA^+^* MAGs were significantly associated with plasma NfL (**Fig. 3d**) or CSF NfL (Extended Data **Fig. 7c**), and *Limosilactobacillus fermentum* was negatively associated with plasma GFAP (**Fig. 3d**). Collectively, these results suggest that *urdA^+^* MAGs are associated with AD-related pathology, including CSF biomarkers of neuropathological aggregates (amyloid-β 42/40 ratio, α-synuclein and p-Tau/amyloid-β 42 ratio), early reactive astroglial response (YKL-40), plasma biomarkers of early tau (pTau-217), and cognitive deficits (mPACC3 scores), reflecting microbial associations with early to moderate stages of AD pathology (**Fig. 3**).

### Genetically increased plasma ImP levels are associated with increased AD risk

The results above suggest ImP and select *urdA^+^* taxa are potential mediators of ADRD phenotypes. To understand potential host genetic factors contributing to inter-individual variations in circulating ImP levels, we used a genetics strategy to investigate whether the observed clinical associations of plasma ImP levels represented a causal relationship. Based on the results of a recent metabolomics genome-wide association study (GWAS), only one locus on chromosome 12p13.33 was identified as being significantly associated with plasma ImP levels^28^. As shown by the regional plot in **Fig. 4a**, the lead variant at this locus is rs7969761 (C>T) and localizes to an intronic region of the *SLC6A13* gene. The T allele of rs7969761 is associated with increased ImP levels (β=0.16; P=7.3x10^-18^) and is present at relatively high frequency (43%) among subjects of European ancestry^28^. We therefore leveraged these results to test the hypothesis that rs7969761 would also be associated with risk of AD dementia. Consistent with this notion, data from the most recent and largest GWAS meta-analysis for AD revealed that the T allele of rs7969761 was directionally consistently associated with increased risk of AD dementia (OR=1.02; 95% CI 1.01-1.04; P=2.8x10^-3^)^29^ . As a more formal test of causality, we also carried out a Mendelian randomization analysis using the Wald ratio estimator, which revealed that a 1 standard-deviation increase in genetically predicted normalized ImP levels was associated with greater risk of AD (OR=1.16; 95% CI 1.05–1.29; P = 3.0 x10^-3^) (**Fig. 4b**). We next used expression quantitative trait locus (eQTL) analysis to prioritize positional candidate genes that could underlie association of this locus with ImP levels and risk of AD. Of note, data from the GTEx Project^30^ revealed multiple *cis* eQTLs with rs7969761 for several genes at this locus, including *SLC6A13, KDM5A, and CCDC77*, in tissues that are involved in AD pathogenesis and regulation of circulating metabolites, such as the brain, skeletal muscle, adipose, and liver. Thus, these results collectively provide genetic evidence that higher circulating ImP levels is causally associated with increased risk of AD.

### Chronic ImP supplementation potentiates AD pathology in male mice

To test whether ImP causally contributes to the progression of AD pathology in a controlled experimental setting, we next examined the effects of chronic ImP supplementation in two complementary mouse models that capture key features of AD (**Extended Data Figure. 8**), including amyloid-β aggregation^31^ and tau pathology^32^.

First, we supplemented 5XFAD mice with ImP orally, via drinking water starting at 4 weeks of age, a regimen designed to increase systemic availability to levels observed in humans, as previously described^9,10,13,33^(**Extended Data Figure. 8**). We detected a significant increase in the number of amyloid-β plaques and percent area occupied by them in the retrosplenial cortex of ImP-treated mice compared to their littermate controls consuming regular water (**Fig. 5a &b**). No differences were detected in the plaques’ average fluorescence intensity (**Fig. 5b**). Since amyloid-β plaques are closely associated with glial cells^34^, we examined astroglial reactivity in these mice. Sections stained for the reactive astrocyte marker GFAP showed a significantly higher number of GFAP^+^ astrocytes in amyloid-β signal dense regions of the retrosplenial cortex (**Fig. 5a &b**). Follow-up proteomics analyses on cortical brain lysates revealed 61 proteins that were upregulated and 10 that were downregulated (fold change cut-off at 30%, FDR adj. p<0.05; 2634 unique proteins detected) in ImP-treated mice compared to untreated littermate controls (**Fig. 5c**) (**Extended Data Table 5**). We observed a significant upregulation of reactive astrocytic proteins SLC1A3 and GFAP, axonal neurofilaments light, medium and heavy, suggesting reactive gliosis and compensatory axonal damage response (**Extended Data Table 5**). We also observed a significant downregulation of Annexin A1, a regulator of the AKT/mTOR pathway^35^ in cerebrovascular injury, a pathway previously implicated in ImP-mediated peripheral vascular injury^13,33^. Additionally, levels of amyloid-beta A4 precursor protein-binding family A member 1 (APBA1), a stabilizer of amyloid precursor protein (APP)^36,37^, were reduced, as were those of platelet-derived growth factor receptor-β (PDGFRβ), whose loss is associated with increased BBB permeability^38,39^ (**Fig. 5c**). Together, these changes suggest that ImP disrupts signaling networks essential for cerebrovascular stability and amyloid homeostasis.

**Figure 5.**
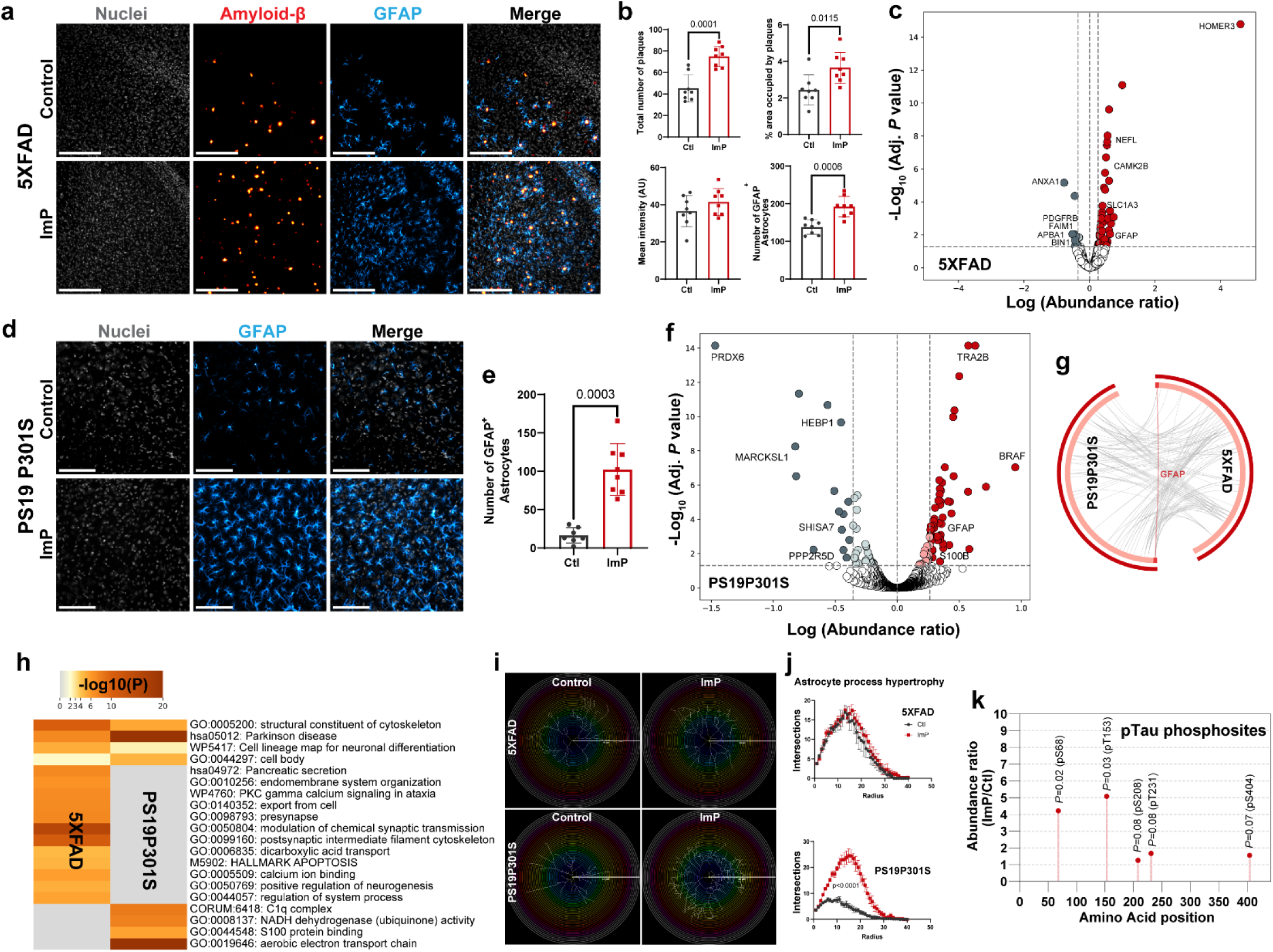
Chronic ImP supplementation exacerbates ADRD-like pathology in male mice. a, Representative immunofluorescence images of amyloid pathology in the retrosplenial cortex of 5XFAD mice (Scale bar= 200µm). **b,** quantification (*N*= 8/group) of amyloid-β plaques (mean ± SD, unpaired t-test), % area occupied by amyloid-β (mean ± SD, unpaired t-test), mean fluorescence intensity of amyloid-β plaques (mean ± SD, unpaired t-test) and total number of GFAP^+^ astrocytes (mean ± SD, Mann-Whitney U-test). **c,** Volcano plot showing differentially abundant cortical proteins in 5XFAD mice (FDR P <0.05, n=4 mice per treatment group; Abundance ratio cut-off 30%). Red and gray dots denote increased and reduced abundances in ImP treated mice respectively. **d,** Representative images of piriform cortex of (*N*= 8) PS19P301S mice (Scale bar= 100µm). **e,** Quantification of GFAP^+^ astrocytes in the piriform cortex of PS19P301S mice (N=8/group, mean ± SD, Mann-Whitney U-test). **f,** Volcano plot showing differentially abundant cortical proteins in PS19P301S mice (FDR P <0.05, n=4 mice per treatment group; Abundance ratio cut-off 30%). Red and gray dots denote increased and reduced abundances in ImP treated mice respectively. **g,** Circos plot illustrating functional overlap (gray) and direct overlap (red) between ImP-induced changes in cortex proteome in 5XFAD and PS19P301S mice, identifying GFAP as a shared differentially abundant protein in both models. **h,** Gene-enrichment analysis of differentially abundant proteins (control *vs.* ImP) in 5XFAD and PS19P301S mice highlighting most significantly enriched pathways in each model. **i,** Representative astrocyte skeletons reconstructions of cortical reactive astrocytes from 5XFAD and PS19P301S mice, overlayed with concentric circles increasing in 1µm increments up to a radius of 40µm (Scale bar= 40µm), centered at the cell body. **j,** Quantification of astrocytes morphological complexity (*N*= 4 mice) using Sholl analysis in cortical reactive astrocytes from 5XFAD and PS19P301S mice (mean ± SEM, repeated measures two-way ANOVA). **k,** Phosphoproteomic analysis of cortical lysates from PS19P301S. ImP treatment increased hyperphosphorylation of ADRD relevant residues S68 and T153 significantly, with increased trends in S208, T231, and S404 within (N= 4, abundance ratio ImP *vs.* control, unpaired t-test & unpaired t-test with Welch’s correction). X-axis represents amino-acid residue numbering mapped to 2N4R h-Tau peptide sequence.

**Figure 6.**
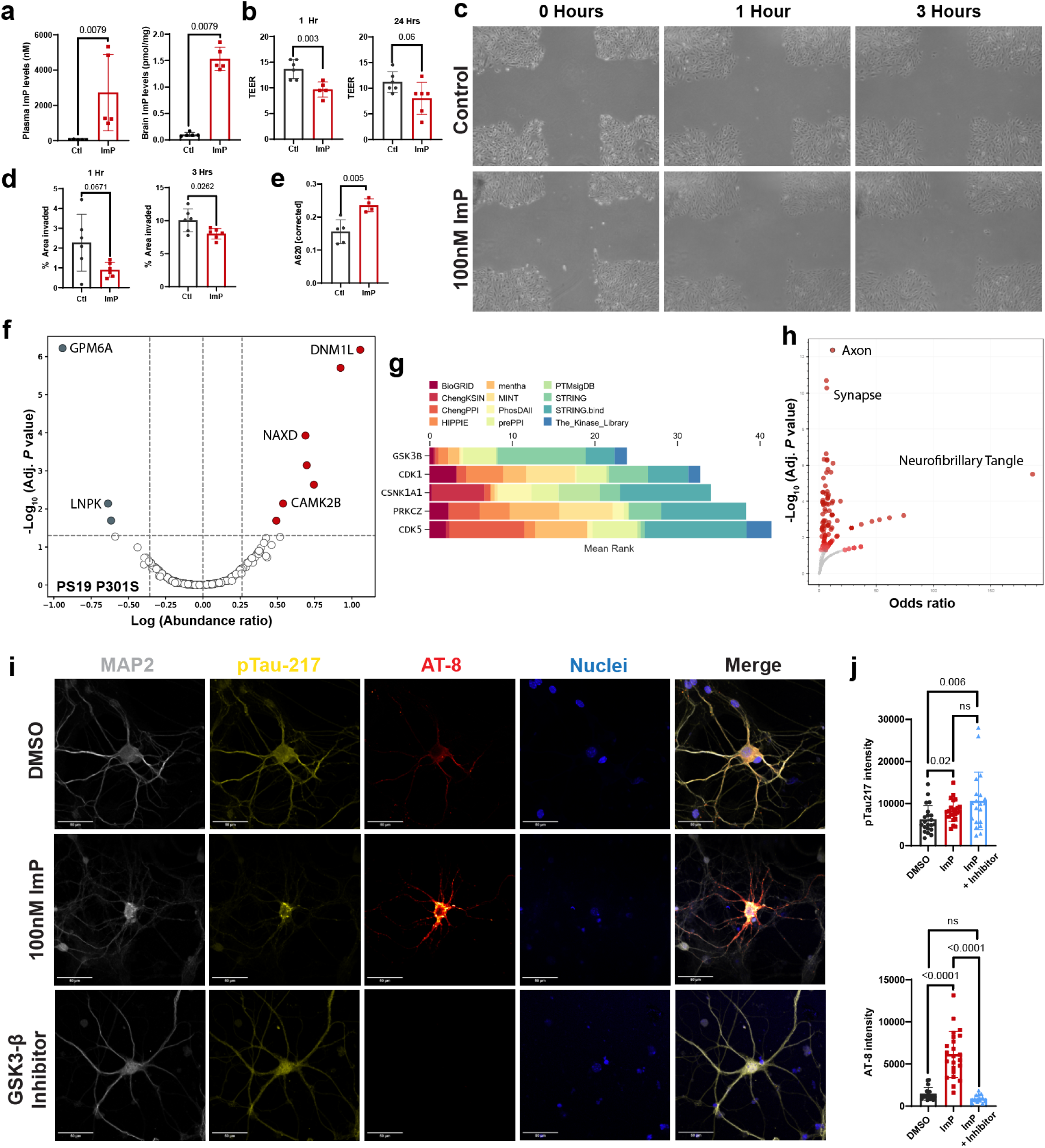
ImP disrupts cerebrovascular integrity and promotes tau hyperphosphorylation through GSK3β. a, Oral ImP administration increases plasma and brain cortical ImP levels in in wild-type C57Bl/6 mice (N=5 per group, mean ± SD, Mann-Whitney U-test). **b,** Treatment of primary human brain endothelial cells (PHBEC) with 500nM ImP reduces trans-epithelial electrical resistance after 1 h with a persistent trend after 24 h (mean ± SD, Mann-Whitney U-test and unpaired t-test). **c,** Representative bright-field images from endothelial wound-healing assay at 0, 1 and 3 h following ImP treatment. **d,** Quantification of PHBEC wound closure following 500nM ImP treatment (*N*= 5 replicates (mean ± SD, unpaired t-test with Welch’s correction). **e,** Oral ImP treatment increases Evans blue extravasation in wild-type C57Bl/6 mice (N = 4 per group). **f,** Phosphoproteomic analysis of cortical lysates from PS19P301S mice following chronic ImP exposure (FDR P <0.05, n=4 mice per group; abundance ratio cut-off 30%). **g,** Kinase enrichment analysis of the top 25 differentially abundant phosphoproteins from PS19P301S cortical lysates, shown as stacked bar plots for top 5 highest-ranked kinases. **h,** Subcellular localization enrichment analysis of top 25 ImP-responsive phosphoproteins from PS19P301S mouse cortical lysates represented as volcano plot with adjusted P values and odds ratios for enriched cellular compartments. **i,** Representative immunofluorescence images of primary cortical neurons treated with vehicle (DMSO), 100nM ImP or 100nM ImP + GSK3β inhibitor. **j,** Quantification of neuronal pTau-217 and pTau-AT-8 (pSer202/Thr205) immunofluorescence (N= 20 neurons/condition, mean ± SEM, Kruskal-Wallis test).

Next, we examined the effect of ImP on tau pathology. We supplemented PS19P301S mice with ImP in drinking water starting at 8 weeks of age. Brains were examined at 6 months of age as previous work suggest that cortical gliosis and tau accumulation are observed at this age^40^. Mice treated with ImP showed significantly higher astroglial reactivity within the piriform cortex, which is in stark contrast with the sparse populations of GFAP^+^ glia detected in age-matched littermate controls (*P* = 0.0003) (**Fig. 5d &e**). Proteomics analysis in cortices identified 2352 unique proteins with 79 that were upregulated and 19 that were downregulated (fold change cut-off at 30%, FDR adj. p<0.05) in ImP-treated mice compared to untreated littermate controls (**Fig. 5f**) (**Extended Data Table 6**). Several proteins associated with reactive astrocytes such as GFAP, S100a1, S100a6 and S100b had increased abundances in ImP-treated mice compared to untreated controls (**Extended Data Table 6**), corroborating histopathological findings (**Fig. 5d**).

Gene enrichment analysis of differentially abundant proteins in both AD mouse models revealed a partial functional overlap (**Fig. 5g &h**). Interestingly, we observe GFAP as the only protein upregulated by ImP, in both 5XFAD and PS19P301S mice (**Fig. 5g**). However, the reactive GFAP^+^ glia detected in both groups of mice appeared morphologically different (**Fig. 5 a, d)**. Thus, we used Sholl analysis^41^ to assess morphology in more detail. We observed that astrocytes from ImP-treated PS19P301S mice expressed increased process-hypertrophy, indicating a more aggressive neuroimmune environment (**Fig. 5i &j)**. This phenotype was not observed in 5XFAD mice (**Fig. 5i &j**), possibly due to glial reactivity being limited proximally to amyloid-β plaques, which are known to perturb homeostatic glia, contributing to their reactivity equally in both control and treated mice^34,42^, but also suggesting a potentially more exacerbated, direct mechanistic link in mouse model of tau pathology.

Lastly, we compared the changes in p-Tau levels at different disease relevant phosphosites in PS19 mice through phosphoproteomic analysis to study pathological hyperphosphorylation of Tau, mapped 2N4R isoform (441 amino acids) for ease of disease stage interpretability. Several ADRD relevant phosphosites were upregulated in ImP treated mice, such as S68, T151, S208 (AT-8), T231 (AT-180) and S404 (**Fig. 5k**), suggesting that chronic ImP treatment increased Tau hyperphosphorylation in 6-month-old male PS19P301S mice, a phenotype that is typically observed at more advanced stages of AD-like pathology^43,44^.

### Imidazole propionate disrupts barrier function of endothelial cells

The results presented above provide experimental evidence linking ImP to ADRD-related phenotypes in mice. We reasoned that any direct effects of ImP on cortical parenchymal cells would require access of ImP to the brain parenchyma, implicating blood-brain barrier permeability as a key step. To test this, we treated one-month-old C57BL/6 wild-type mice with ImP for two months (**Extended Data Fig. 8**). As expected, ImP-treated mice showed significantly elevated plasma ImP levels (**Fig. 6a**). Analysis of cortical tissue lysates following transcardiac perfusion with saline (to remove blood) revealed marked accumulation of ImP within brain tissue (**Fig. 6a**), demonstrating that ImP penetrates into the cortex and may directly interact with neurons and glial cells. This was surprising because *in-silico* analysis of ImP physicochemical properties^45,46^ suggested that although it is likely absorbed in the intestines, it is not predicted to freely cross the BBB (**Extended Data Table 7**). Because ImP has been shown to induce vascular endothelial injury in peripheral tissues^13,33^, we investigated its effects on primary human brain microvascular endothelial cells. Exposure of endothelial monolayers to ImP significantly reduced transepithelial electrical resistance (TEER) (**Fig. 6b**), indicating loss of barrier integrity. ImP also impaired wound-healing capacity (**Fig. 6c & d**), consistent with diminished endothelial repair activity and previous reports in aortic and umbilical vein endothelial cells^13,33^. Given that brain endothelial cells interact closely with astrocytes and pericytes, which help preserve cerebrovascular integrity^47–49^, we next assessed vascular permeability *in vivo* in ImP-treated C57BL/6 mice using Evans blue dye. Brain lysates from treated mice showed significantly increased absorbance at 620 nm (**Fig. 6e**), indicating enhanced dye extravasation into cortical tissue. Together, these findings demonstrate that ImP compromises endothelial barrier function and promotes vascular leakage within the brain.

### Imidazole propionate promotes tau hyperphosphorylation through GSK3β

The results above show that ImP accumulates at significant levels in brain tissue (**Fig. 6a**), raising the possibility that it could directly interact with neurons. ImP was previously linked to AKT activation through p38γ^13,33,50^ in peripheral endothelial, myeloid and human embryonic kidney (HEK293) cells, a signaling axis that in neurons can drive tau hyperphosphorylation^51,52^, axonal destabilization^53^, and trigger reactive astrogliosis. To study potential alterations in kinase activity, we performed untargeted analysis for alterations in cortical phosphoproteome of ImP treated PS19P301S mice. We detected 356 unique phosphorylated proteins with 7 upregulated and 2 downregulated (fold change cut-off at 30%, FDR adj. p<0.05) phospho-sites in ImP-treated mice compared to untreated littermate controls (**Fig. 6f**) (**Extended Data Table 8**). We observed increased phosphorylation of axonal proteins such as microtubule associated proteins 1A, 1B, and tau; tubulin alpha and beta; neurofilaments light and heavy etc. Because kinase signaling involves complex multi-step cascades, with overlapping upstream activators and downstream targets, we used KEA3 (Kinase enrichment analysis)^54^ to infer and rank kinases potentially involved in phosphorylating the top 25 phosphopeptides showing the largest change in our dataset. This analysis identified GSK3β as the top candidate, with the best mean rank score across all databases and mapping to 14 target proteins (**Fig. 6g**). In comparison, previously reported interactors for ImP, namely, p38γ (mean rank score >100; 7 target proteins), AKT (mean rank score = 19; 17 target proteins) and mTOR (mean rank score = 26; 13 target proteins) (**Extended Data Fig. 9**) showed lower overall enrichment. Since cortical tissue has heterogeneous populations of neurons, glia, endothelial cells etc., we enriched for cellular compartments^55^ from the phosphoproteome to identify specific cells and their sub-cellular compartments that are affected by ImP, revealing axon, synapse and neurofibrillary tangles as the strongest result, suggesting that ImP’s affects are most significant in compartments within neurons (**Fig. 6h**).

The observed increase in kinase activity *in vivo*, in neuronal cytoskeleton may reflect a direct effect of ImP or arise secondarily from cellular stress, neuronal damage, and glial reactivity. To distinguish between these possibilities, we treated primary neurons from PS19P301S mice, seeded with 2N4R h-Tau pre-formed fibrils, with 100nM ImP. Because GSK3β is a central kinase and an early mediator of tau hyperphosphorylation^56^, and was implicated in ImP-driven signaling by our phosphoproteomic analysis (Fig. 6g), we also co-treated neurons with the GSK3β inhibitor, N-(4-Methoxybenzyl)-Nʹ-(5-nitro-1,3-thiazol-2-yl) urea, a known tau phosphorylation inhibitor^57^. ImP treated neurons showed more than 3-fold increase in pTau-AT-8, relative to controls (**Fig. 6i & j**). This effect was fully reversed by GSK3β inhibition (**Fig. 6i & j**). These results suggest that ImP promotes tau phosphorylation, at least in part through GSK3β.

## DISCUSSION

The findings presented here identify the gut bacterial metabolite ImP as a potential early modifier of AD pathogenesis, shaped by host genetic variation and acting through both neurodegenerative and cerebrovascular mechanisms. Previous studies have established that gut microbiome alterations occur in individuals with AD^3–5^. This study advances this concept by linking a specific gut-derived metabolite to clinical and molecular features of disease. Elevated plasma ImP concentrations were associated with multiple biomarkers of AD pathology, neurodegeneration, and cognitive decline in a large, predominantly cognitively unimpaired cohort. Importantly, complementary genetics and experimental data in humans and preclinical models, respectively, demonstrated that elevated ImP levels were causally associated with AD-relevant phenotypes, highlighting the pathogenic potential of this gut microbiome-derived metabolite. These findings thus support the notion that products of microbial metabolism of dietary nutrients can influence brain function of the host well before the onset of clinical symptoms.

This work extends prior studies by showing that inter-individual variation in circulating ImP reflects both microbial and host determinants, including age, sex, and genetics. Prior work identified *E. lenta*, *S. wadsworthensis*, and *A. equolifaciens* as ImP producers in metabolic disease contexts^9,10,12,27^, and our metagenomic analysis supports these observations while broadening the list of candidate ImP-producing taxa harboring putative *urdA* homologs (**Extended Data Fig. 4a**) and associated with AD biomarkers. Additionally, environmental inputs, particularly diet, likely contribute to this heterogeneity by altering the abundance and activity of ImP-producing bacteria, providing one route by which microbiome composition may shape systemic ImP levels^9,33^. A key advance is that host genetics appears to contribute to ImP regulation and, in turn, to AD susceptibility, adding a layer beyond microbiome composition and diet. The genetic signal supports a model in which higher ImP lies on a causal path to disease risk. The emergence of a single locus influencing circulating ImP levels, which is also associated with AD risk, suggests a shared host genetic mechanism linking ImP regulation and disease susceptibility. Moreover, eQTL evidence at the locus prioritizes several positional candidates with plausible roles in metabolite handling and AD-relevant biology across multiple tissues (including brain and metabolic organs), raising the possibility that genetically driven differences in transport, cellular uptake, or systemic clearance may impact circulating ImP levels. Together, these results indicate that circulating ImP levels are shaped by both bacterial (e.g., *urdA* carriage and activity) and host factors (age, sex, and genetics), with downstream consequences for neurovascular integrity and neurodegeneration. They also point to the need to identify the causal gene(s) at the chromosome 12 locus and underlying mechanism for ImP regulation.

Using two complementary mouse models of AD, we further show that elevating ImP levels through chronic dietary exposure worsened key features of AD pathology. In the amyloid pathology model, we observed an increased deposition of amyloid-β plaques within the cortex, with a concomitant increase in glial reactivity and blood-brain barrier damage markers (**Fig. 5**). In the tau model, these effects were accompanied by significant changes in the brain proteome and phosphoproteome, including increased abundance of reactive astrocyte markers and elevated phosphorylation of tau at disease-relevant residues. Mechanistically, the phosphoproteomic data indicates increased activity of kinases that regulate phosphorylation of axonal cytoskeletal proteins. Altered phosphorylation of neurofilaments and microtubule-associated proteins is known to compromise axonal structure, disrupt axonal transport, and lead to neuronal dysfunction and loss, which are central to AD pathology^58^. The protein transformer-2 protein homolog β (TRA2B), which is a known regulator of exon 10 splicing of tau associated with fronto-temporal dementia with Parkinsonism-17 (FTDP-17)^59^ was among the most abundant proteins upregulated, potentially linking ImP to alternative splicing events relevant to tauopathies. This mechanism could potentially be controlled by serine/arginine repetitive matrix protein 1 (SRRM-1), which can bridge splicing machinery to enable TRA2B function in the production of pathogenic Tau^60^. Furthermore, ImP-treated mice expressed higher levels of phosphorylated dynamin-1-like protein (DRP1) (**Extended Data Table 7**), which is involved in mitochondrial fission^61^, potentially contributing to upregulation of mitochondrial reactive oxygen stress markers in ImP treated mice (**Extended Data Table 7**). Together, these findings suggest that ImP not only amplifies established features of AD pathology but may also engage broader neurodegenerative processes, including glial activation, dysregulated tau phosphorylation and handling, and cellular stress responses that contribute to disease progression.

The ImP-driven increase in amyloid pathology in 5XFAD mice, may be partly explained by vascular stress signatures detected by histology and proteomics (for example, GFAP, SLC1A3, and PDGFRβ), consistent with impaired perivascular/glymphatic clearance. By contrast, PS19 P301S mice showed a profile, characterized primarily by changes in axonal cytoskeletal proteins (**Extended Data Tables 6 & 8**), consistent with the phosphoproteomic signatures described above. Kinase enrichment analysis of the differentially phosphorylated proteins identified GSK3β as a central node—a stress-responsive kinase in ADRD that phosphorylates disease-relevant residues in MAPT. In line with this, our in vitro data show that GSK3β inhibition reverses ImP-induced AT8 phosphorylation, a tau modification linked to cytoskeletal disruption in later stages of disease. Furthermore, pTau-217 observed in humans and in the neuronal model is phosphorylated by GSK3β or CDK5, the kinetics of these interactions should be investigated in future studies. Kinase inhibition may therefore be relevant for individuals with high ImP and increased vulnerability to tauopathy. However, upstream approaches, such as reducing ImP production in the gut or limiting its entry into the circulation, may offer broader protection, especially given associations between ImP and type 2 diabetes, atherosclerosis, and kidney disease.

This study has limitations which should be mentioned. Our human cohort consisted primarily of cognitively unimpaired individuals, which may limit the generalizability of our findings to broader or more diverse populations, including those with varying degrees of cognitive impairment or differing geographic and demographic backgrounds. However, two recent large studies have linked ImP with cognitive decline, including the Medical Research Council (MRC) National Survey of Health of Development (NSHD) cohort (*N* = 1,740)^62^ and the Rotterdam study (*N* = 1,082)^63^, with results matching our findings (**Fig. 1**). While the cross-sectional design for the clinical associations precluded assessment of temporal or causal relationships in humans, Mendelian randomization analyses with a limited number of instrumental variables did provide evidence that the relationship between ImP and cognitive traits in humans was causal. However, longitudinal studies will still be needed to determine whether increases in microbiome-derived metabolites, such as ImP, contribute to longitudinal cognitive decline over time, although these are difficult to perform due to the extended follow-up period needed. To partially address this limitation, we modeled trajectories of cognitive performance and plasma biomarkers over age in individuals stratified by ImP status (**Fig. 1f & 2b**). These predictions are based on statistical assumptions and require validation in longitudinal cohorts. ImP measurements for all of the associations were also acquired by untargeted metabolomics with arbitrary values by abundance, that were harmonized and median normalized. Since these measurements do not provide molar concentrations, it will also be important to quantitate absolute levels of ImP to derive more accurate effect sizes for the associations observed in humans and mice ImP was administered orally in our preclinical studies. While this controlled experimental approach effectively elevates plasma ImP levels^13,33^ and likely minimized variation in humans that could have be introduced by diverse geographical, lifestyle, comorbid, dietary, and genomic patterns, oral delivery *vs.* gut production may alter ImP kinetics and host responses. Future validation using gnotobiotic models colonized with defined microbial communities differing in ImP-producing capacity will be essential, though such models will still require optimization. Furthermore, in the 5XFAD model, which is sensitive to insulin signaling^64^, the known effects of ImP on insulin signaling^10^ may have contributed to the observed glial phenotype. It has also been recently discovered that paternal inheritance of disease transgenes in 5XFAD mice may determine disease severity in offsprings^65^. In our study, however, littermate controls were used consistently, minimizing any potential confounding effects of parental genotype on the observed phenotypes.

Finally, while ImP-induced tau phosphorylation in neurons was significantly reduced by GSK3β inhibition, further work is needed to define the precise signaling hierarchy linking ImP to tau pathology. Given that ImP has been associated with AKT signaling in peripheral tissues^13,33,50^ and that AKT inhibits GSK3β via phosphorylation^66^, future experiments should address the temporal and dose-dependent relationships among ImP, AKT, and GSK3β. To prioritize kinases potentially mediating ImP’s effects, we employed a mean rank-based scoring approach across multiple databases and identified GSK3β exhibiting greater predicted downstream activity than AKT or mTOR in cortical tissue from 6-month-old PS19 P301S mice. Importantly, a recent study of ImP in Parkinson’s disease reported differential activation of mTOR, but not AKT^67^, whereas another study examining stress-induced behaviors found no change in mTORC1 activation^68^. The pathological processes altered by ImP were limited to astrocytes but no differential signal is observed in mice for microglial activation other than C1Q and C2Q in PS19P301S mice, this likely implies that ImP’s effects in our study show bias to astroglia, the recent study exploring stress-induced behaviors under ImP similarly reported no changes in microglia even under isolated conditions^68^. Collectively, these observations suggest that ImP’s effects in the brain are pleiotropic and may vary depending on disease context and tissue. Further studies are warranted to elucidate the molecular mechanisms linking ImP to AD pathophysiology, including its potential receptors or transporters, as well as to evaluate whether targeting upstream ImP production or specific downstream signaling pathways may offer therapeutic benefit.

Despite these limitations, the results described here present a remarkably consistent pattern where elevated ImP impacts multiple indices of AD pathology, including amyloid-β, phosphorylated tau, neurodegeneration, glucose metabolism, and cognition among humans, and measures of AD and cerebrovascular pathology in murine models (**Extended Data Fig. 10**). These results open new avenues for microbiome-targeted interventions, such as modulation of ImP-producing taxa^69^, inhibition of UrdA enzymatic activity, dietary strategies to limit histidine-to-ImP conversion^70^, or inhibition of ImP uptake into systemic circulation. As gut-brain axis research advances, identifying and targeting specific microbial metabolic routes, like ImP production, may provide novel opportunities for early prevention or slowing of AD progression.

## MATERIALS AND METHODS

### Study design

The overall objective of this study is to test whether the gut-bacterial metabolite ImP is a potential risk factor for exacerbated disease outcomes in ADRD. Human subjects’ data was obtained from WADRC, WRAP, and MARS cohorts, all of which are observational studies. The procedures for phenotype measurements are fully described in the corresponding Methods sections.

Experiments featuring transgenic pre-clinical murine models of AD are comprised of heterozygous transgenic mouse litters split into equal/nearly equal numbers of control and treatment groups. A final *N* = 4 was achieved per cage, with two different cages in each treatment group for a total of *N* = 8 mice per treatment group as two individual experiments. For PS19P301S mice, *N* = 4 mice/group were bred locally (using breeders obtained from Jackson Labs), whereas the remaining 4 mice/group were purchased from Jackson Labs separately. Effect sizes were determined from previous studies to obtain an approximate N for the experiments. Mice were exposed to ImP chronically, via drinking water. One data point was excluded from **Fig. 6e** due to an abnormal amount of dye pooling at tissue level, post-mortem. The number of replicates used in histopathological analysis was mentioned in associated methods. All images were blinded using an in-house bash script, unblinded only for plotting the resultant data and obtaining representative images.

### Participants

Participants were recruited from the ADRC and WRAP studies^19^. The Wisconsin ADRC clinical core recruits individuals representing the full clinical and biological spectrum of AD, ranging from cognitively unimpaired individuals to those with mild cognitive impairment and AD dementia. The WRAP is a longitudinal observational study that enrolls participants from a cohort enriched with individuals at increased risk for late-onset AD due to a parental history of AD dementia. Participants underwent *APOE* genotyping using competitive allele-specific PCR-based KASP™ assays (LGC Genomics, Beverly, MA) and participated in longitudinal assessments of cognitive function and laboratory tests. A subset of the cohort provided CSF samples collected via lumbar puncture and underwent PET neuroimaging to assess biomarkers of AD and related pathologies. Additionally, participants co-enrolled in the Microbiome in Alzheimer’s Risk Study (MARS) provided fecal samples for gut microbiome analysis and completed questionnaires related to medical history and diet at the time of sample collection. Participants provided informed consent following the guidelines of the Declaration of Helsinki prior to enrollment in the Wisconsin ADRC, WRAP, and MARS studies. The study procedures were approved by the Institutional Review Board of the University of Wisconsin (Wisconsin ADRC and WRAP umbrella protocol ID: 2013-0178; MARS protocol ID: 2015-1121). Previous studies reported measurements of plasma ImP, mPACC3 scores, CSF biomarkers, and FDG PET, and further methodological details are available in Supplementary Materials and Methods.

### Metagenome Assembled Genomes and *urdA* gene annotation

Shotgun sequencing reads were assembled for all samples using SPAdes (v.3.15.5; metaspades.py -k 21,33,55,77), the assembled contigs were quantified by mapping shotgun reads to contigs using Bowtie2 (v.2.3.4). Contigs that were less than 500 bp were excluded for downstream analyses. Contigs were then binned into MAGs using MetaBAT2 (v.2.17). The quality of all MAGs was assessed by genome completeness and contamination using CheckM (v.1.2.2) and high-quality MAGs were kept (completeness > 90% and contamination < 5%). The high-quality MAGs were dereplicated using dRep (v.3.5.0; -pa 0.9 - sa 0.99). The final dataset contains 1464 MAGs. Taxonomic assignments of these 1464 MAGs were using the Genome Taxonomy Database Toolkit (GTDB-Tk; v.2.3.2) and the GTDB database (ver. R214). Genes from MAGs were predicted using Prodigal (v.2.6.3) and annotated to KEGG Orthology database using kofamscan (v.1.3.0) and annotated to Carbohydrate-Active Enzymes (CAZymes) database using dbcan3 (v.4.1.4) and database “dbCAN3_db_v12_20240415”. MAGs were processed to create a single kallisto quantification index and reads from each fecal DNA sample were mapped to this index to quantify the abundance of each MAG in each sample^71^.

Hidden Markov model (HMM) was built using *urdA* reference genes from a collection of known ImP-producing bacteria^10^. Reference *urdA* gene sequences were aligned using MUSCLE (v.3.8.31)^72^, then the HMM profile was constructed using “hmmbuild” from HMMER (v.3.2.1). Protein coding genes (CDS) from MAGs were searched using HMM search (v.3.2.1). All genes involved exhibiting 60% score of the lowest scoring model sequence were included in subsequent analysis. Clustal Omega^73^ was used for multiple alignment of predicted UrdA genes, and potential UrdA homologs (producing imidazole propionate from urocanate) were defined by “Y” or “M” residue at the FAD active sites tyrosine 373 (Y373).

### Animal husbandry

All animals were obtained from The Jackson Laboratory, after which they were housed and bred in a specific-pathogen-free (SPF) facility at the University of Wisconsin-Madison. Male mice were split into cages of 4 with littermates evenly divided between test and control groups, and water intake was carefully monitored to obtain the average amount of water consumed per mouse within a cage for all cages. Imidazole propionate (Cayman Chemical) stock solution was prepared at 200µg/mL to match the average daily intake of water of mice at 4mL/day to enable a dosage of 800µg/mouse/day. Wild-type C57BL/6J mice were treated for 4 weeks before Evans blue extravasation assay and harvest. 5XFAD [B6.Cg-Tg(APPSwFlLon,PSEN1*M146L*L286V)6799Vas/Mmjax), Tg6799] mice were supplemented with ImP starting at 2 weeks of age until they were 6 months-old, whereas PS19P301S [B6;C3-Tg(Prnp-MAPT*P301S)PS19Vle/J] mice were treated starting at 8 weeks of age until they were 6 months-old. All protocols for this study were approved by the Institutional Animal Care and Use Committee (IACUC) at the University of Wisconsin-Madison.

### Mouse brain sample collection and confocal microscopy

Mice were anesthetized with isoflurane [Isospire, Dechra] and blood was collected from the right ventricle. Mice were then perfused transcardiacally through the left ventricle with 15mL perfusion buffer consisting of PBS and heparin (0.5 USP). Brains were bisected into left and right hemispheres, where the left hemisphere was flash frozen and right hemisphere was fixed in 4% PFA for 48 hrs at 4C followed by fixation in 30% sucrose until the brains reach equal relative density to the sucrose solution. The brains were then fixed in sucrose [Fisher scientific] -OCT [Tissue-Tek, Sakura Finetek] solution for sectioning. Brains were sectioned into 40µm sections from anterior to posterior and placed into an anti-freeze solution comprised of PBS [VWR Life science], sucrose and ethylene glycol [Sigma- Aldrich]. Sections from morphologically comparable regions with visible hippocampus, retrosplenial cortex and piriform cortex located 240µm apart from each other, were then washed with PBS and incubated with antibody for reactive astrocytes [GFAP (clone: GA5) 488 Conjugated, Invitrogen, 1:1000] for 12 hrs in 4C followed by another PBS wash and incubation with stain for amyloid-β [MethOxy XO4, Tocris, 1:1000] and nuclei [TO-PRO-3 Iodide (642/661), ThermoFisher Scientific, 1:1000] at room temperature for 1 hour. The sections were then washed and mounted using a mounting solution [Fluoromount G, Invitrogen]. 3 images were taken from random regions within retrosplenial cortex in 5XFAD mice and piriform cortex in PS19P301S mice, from 3 different coronal sections using Zeiss LSM 800 (UW-Madison, Medical Microbiology and Immunology) and Nikon A1R microscopes (UW-Madison Biochemistry Optical Imaging Core).

### Image analysis

Immunofluorescence data was quantified with the help of the surfaces and spots function on Imaris (Oxford Instruments). To count the number of GFAP+ cells, spots were generated with 6µm diameter localized to the center of astrocyte soma. Generated spots were cross-checked with spots generated by co-localizing nuclei with GFAP to obtain nuclei within GFAP+ areas as a proxy for cells. Surfaces function was used to render 3D solids using the amyloid-beta channel at fixed thresholds over all images, to render amyloid-beta plaques. A specialized MATLAB [Mathworks] script was used to separate individual plaques to obtain information on average volume, area and intensity. To quantify the hypertrophy in astrocyte processes, images were rendered on Image J (National Institutes of Health) where, 40µm thick hyper-stacks were converted into a 2D Z-project. A binary image was generated using the threshold feature, a region of interest was drawn around an isolated astrocyte and data outside the ROI was cleared. The skeletonize function was used to generate an astrocyte skeleton and we used SNT neuroanatomy plugin^74^ set at 1µm intervals for Sholl analysis from the center of the skeleton up to 40µm away from the center. Image J’s threshold function was utilized to measure the area fraction of MethoxyX04 positive signal. Neuronal pTau and MAP2 signals were quantified as mean intensities within ROI drawn encapsulating the neuronal soma of healthy, isolated neurons.

### Proteomics and phosphoproteomics

Proteins were extracted from flash-frozen cortices of PS19P301S male mice, followed by methanol-chloroform precipitation, tryptic/LysC digestion, and TMT labeling, as previously described (see Supplemental Methods). Phosphopeptides were enriched using sequential TiO₂ and Zr-IMAC magnetic bead-based protocols and analyzed by nanoLC-MS/MS on an Orbitrap Fusion Lumos platform. Details provided in Supplemental Methods.

### Measurement of ImP *in vivo*

Plasma samples were obtained via cardiac puncture. Brain tissue samples were collected following cardiac puncture and transcardiac perfusion with PBS, to clear blood from the cortical tissue, followed by immediate flash freezing. Cortex was dissected from the flash frozen brain and homogenized for ImP measurement as described in the references^33,67^.

### Evans blue extravasation assay

Mice were anesthetized as mentioned above, 1% Evans blue solution [Thermo Scientific] was prepared and filtered through 2µm syringe filter [Millipore]. 150uL of Evans blue solution was injected retro-orbitally and mice were allowed to recover for 30 minutes. Mice were anesthetized and perfused transcardiacally with perfusion buffer as mentioned above. Brains were collected, flash frozen and homogenized using a bead-beater with 0.1 mm zirconia beads [Biospec]. Homogenates were centrifuged at 10,000 RPM for 20 minutes at 4C, supernatant was collected, and absorbance was measured at 620nm. Final values were normalized by subtracting OD620 of a mouse brain without Evans blue treatment.

### Cell culture assays

Human brain microvascular endothelial cells (HBMVEC) were purchased from iXCells Biotechnologies and grown in Endothelial Cell Growth Medium 2 and the corresponding supplements [PromoCell]. Cells were cultured in 5% CO2 at 37°C. Experiments were performed when cells reached confluency. HBMVEC were used between passage 4 and 6. Cells were treated with Imidazole Propionate at 500 nM through the growth medium. To evaluate wound healing, cells were cultured under normal conditions as above. At confluency, the monolayer was scratched with a P200 pipette tip in a plus (+) shape within the well and washed with PBS. The cells were imaged at 15 minutes, 3 hours and 24 hours. Images were analyzed on Image J by measuring percentage area invaded into the scratched region of interest. To test the membrane integrity, cells were cultured in transwell plates [Corning] with 500nM ImP within medium. TEER was measured using Electrical Resistance System and STX electrode [Millicell]. Final values were corrected for resistance of the blank membrane.

Primary cortical neurons were isolated from the brains of P0 PS19P301S mice. Briefly, the cortices were harvested from pups into ice-cold HBSS (Hank’s balanced salt solution), where the midbrain and meninges were removed. Brain tissue was minced and digested in 0.25% trypsin for 20 minutes at 37°C. Trypsin was quenched with DMEM (Dulbecco’s Modified Eagle Medium)/10%FBS (fetal bovine serum)/1%Penicillin/Streptomycin, and cells were dissociated and counted prior to plating on poly-d-lysine coated glass coverslips. Cultures were then fully changed to Neurobasal Plus Media (2% B27+, 1% GlutaMax, 1% Penicillin/Streptomycin) the subsequent day. Upon the first change to Neurobasal media, neurons were treated with 1μM AraC (cytosine arabinoside) to eliminate glial cell populations. The media was changed by ½ volume every 3–4 days until the experiments. To induce neurofibrillary tau tangle accumulation in the P301S, transgenic neurons were seeded with 3mg/mL of pre-formed fibrils (PFF) [SPR-329; StressMarq] on day 7 in vitro and then cultured for 12 days. At day 12 in vitro, primary neurons were treated with 100nM ImP for 96 hours. Media was changed after 72 hours and replenished with media containing100nM ImP.

After 96 hours of treatment, cells were fixed in 10% formalin for 10 minutes, permeabilized with 0.3% Triton X-100 in PBS for 1 hour and incubated with a primary antibody cocktail overnight. Primary antibodies used: MAP2 [1:5000 dilution; ab5392; Abcam], AT-8 [1:500 dilution; MN1020; Thermo Scientific], and pTau Ser217 [1:250 dilution; 44-744; Thermo Scientific].

### Statistical analysis

All statistical analysis on the human data was performed with Python (v3.12.1). The age difference between sample collections from two datasets was calculated for any dataset combined with plasma ImP data, and samples with an age difference greater than 4 years were filtered out. The Mann–Whitney *U* test was used to analyze differences in ImP levels between females and males. Spearman correlation was performed to examine the relationship between age and ImP levels. These statistical analyses were performed with SciPy (v1.11.4). To test the relationship between ImP and cognitive performance, the ordinary least squares (OLS) multiple linear regression analysis was performed accounting for age, sex, BMI, *APOE* ε4 carrier status, and the age gap between plasma sample collection and assessment of cognitive performance using statsmodels (v0.14.1). To reduce the impact of highly influential observations, Cook’s distance was calculated for each OLS model, and observations exceeding the standard cutoff of 4 / (n − p) were removed before refitting. The association between ImP and plasma biomarkers, including NfL and pTau-217, was examined with the OLS regression model controlling for age, sex, BMI, *APOE* ε4 carrier status, and the age gap between plasma samples collection using statsmodels (v0.14.1). The association between ImP and CSF biomarkers with Spearman correlation was performed using SciPy (v1.11.4). The relationship between ImP and FDG PET SUVR of the frontal, temporal, and parietal lobes was determined with the OLS regression model using statsmodels (v0.14.1) adjusting for covariates including age, sex, BMI, *APOE* ε4 carrier status, and the age gap between plasma sample collection and FDG PET imaging. For the longitudinal analysis, participants were first stratified into one of four groups—Low, Medium-low, Medium-high, or High—based on the quartiles of their average ImP values. Group assignment was performed using equal-frequency binning via the pd.qcut function (pandas v2.2.1) to ensure approximately equal group sizes. We then filtered for only the Low and High ImP groups to assess differences in longitudinal trajectories. To examine the relationship between the ImP group and projected trajectories of plasma biomarkers or cognitive scores over age, we fit a mixed-effects model using the mixedlm function (statsmodels v0.14.1). The model specified the outcome (cognitive scores or plasma biomarkers) as predicted by centered age (each participant’s age − mean age) interacting with ImP group, along with sex, *APOE* ε4 carrier status, BMI, age difference between samples, and practice effect (included only for repeated cognitive testing), with a random intercept for each participant to account for within-subject correlations. Although the number of observations per subject varied, with some individuals contributing up to three timepoints and others fewer, mixed-effects models inherently accommodate such unbalanced longitudinal designs. This approach allows each available observation to contribute to parameter estimation without requiring matched follow-up between groups. The models were fit using restricted maximum likelihood (REML), which provides unbiased estimates in the presence of unbalanced data, assuming data are missing at random. Multiple test correction was performed using the Benjamini–Hochberg procedure to control for false discovery rate using statsmodels (v0.14.1). All figures were generated using Matplotlib (v3.8.0) and seaborn (v0.13.2).

Multiple linear regression models were used to relate the abundances of MAGs with CSF biomarkers using the formula: *CSF biomarker ∼ MAG + sex + age + BMI + APOE allele + NTK batch + age difference between fecal and CSF samples*. The alignment file generated using Clustal Omega was converted to FASTA format using AlignIO.convert() function from Biopython (v1.85). Phylogenetic analysis was conducted using IQ-TREE with the MFP (Model Finder Plus) model, and bootstrap values were calculated with 1000 replicates (-bb 1000) and approximate likelihood ratio tests (ALRT) with 1000 replicates (-alrt 1000). The optimal number of threads was automatically determined (-nt AUTO). The resulting tree file was subsequently visualized and annotated using the Interactive Tree of Life (iTOL, v7.1) tool^75^.

The TPM (Transcripts per kilobase per million) output of kallisto followed by centered log ratio (CLT) transformation was used as MAG abundances in each sample. The beta coefficient for MAG from the formula was used to determine associations between CSF biomarkers with MAGs. Additionally corrected for batch effects from NTK samples using “*NTK batch”* covariate.

All statistical tests pertaining to mice and in vitro models are performed on GraphPad [PRISM v.10.3.1(509)]. Data was expressed in mean±SD or mean±s.e.m described under figure legends for appropriate graphs. Statistical significance was assessed utilizing two-tailed or one-tailed unpaired Student’s *t*-tests, two-tailed or one-tailed Welch’s *t*-test, Mann-Whitney U test, two-way repeated measures ANOVA, and Kolmogrov-Smirnov test, described per figure basis, decided on distribution of data points and differences standard deviation between test and control groups. The level of significance was set for *P* value<0.05.

Raw data pertaining to proteomics and phosphoproteomics was directly imported into Proteome Discoverer 3.1.0.638 where protein identifications and quantitative reporting were generated. Seaquest HT search engine platform with INFERYS rescoring-based workflow for high-resolution MS2 TMT quantification was used to interrogate Mus musculus (Mouse) reference proteome database (UP000000589, 04/19/2024 download, 54,874 total entries) containing human MAPT protein (P10636 UniProtKB accession). Peptide N-terminal and lysine TMT labeling plus Cysteine carbamidomethylation were selected as static modifications whereas methionine oxidation plus serine and threonine phosphorylation were selected as variable modifications. Peptide mass tolerances were set at 10ppm for MS1 and 0.03Da for MS2. Peptide and protein identifications were accepted under strict 1% FDR cut offs with high confidence XCorr thresholds of 1.9 for z=2 and 2.3 for z=3. For the total protein quantification processing Reporter Ion Quantifier settings were used on unique peptides, protein grouping was considered for uniqueness. Reporter abundance was based on normalized total peptide amount intensity values with co-isolation threshold filter set at ≤50. t-test (background based) hypothesis was executed without imputation mode being executed. For the phospho-enriched analysis no global normalization was applied, and only phosphorylated peptides were allowed for pairwise ratios quantification.

## Supporting information

Figure 3. Panel a

## Acknowledgements

We extend our thanks to the WRAP and Wisconsin ADRC participants for their involvement and making this work possible. We thank the staff and researchers at the University of Wisconsin ADRC, the Wisconsin Alzheimer’s Institute, the Wisconsin Institute for Medical Research, and the Waisman Center for their assistance in study organization, participant recruitment, and data collection, the University of Wisconsin Center for High Throughput Computing (CHTC) in the Department of Computer Sciences for providing computational resources, support and assistance.

## Author contributions

F.E.R., F.B., B.B, V.V. conceived the study. V.V., J.W.K., B.B. and F.E.R. designed experiments. V.V. led the study and performed all mouse experiments. J.W.K., Y.D., R.S., analyzed clinical data. H.B., J.F.K., S. C. J., S. A., H. Z., K. B., C.D.E., R. M. A., T. K. U., F.E.R., B. B. B. contributed valuable resources and datasets. Q.Z., J.W.K., performed metagenomic analysis. S.H., J.L.H coordinated sample collection. K.R.B performed metabolite measurements. V.V., E. R. M., R.A-M performed cell culture studies. J.H., H.A. performed GWAS analysis and MR analysis V.V., J.W.K and F.E.R. wrote the manuscript. All authors approved the final manuscript.

## Data and materials availability

All clinical data are available in previous publications mentioned in the respective methods section. The metagenomics dataset included in this study was deposited in the Sequence Read Archive (SRA) under accession number PRJNA1271016.

## Supplementary Materials and Methods

### Plasma sample collection and ImP analysis

Fasting blood samples were drawn from study participants from the WRAP study as previously described^1,2^. Plasma samples were obtained from the blood samples by centrifugation at 3000 revolutions per minute for 15 minutes at room temperature. Plasma levels of ImP were determined by HPLC/MS (Metabolon, Inc.), as previously described^3^. The raw metabolite data was median scaled, log-transformed, and K-Nearest Neighbors (KNN) imputation was applied to fill in any missing metabolite values using the KNNImputer from the sklearn.impute module (v1.4.1), with five nearest neighbors. The imputed ImP data was used for downstream analyses.

### Plasma biomarkers pTau-217 and NfL

The commercial ALZpath pTau-217 assay was used on the Simoa^®^ platform to analyze plasma pTau-217 levels from WRAP participants. Plasma biomarkers, including NfL, from WRAP were analyzed and quantified using the commercial Neurology 4-plex E kit (103670; Quanterix). All assays were performed at the Department of Psychiatry and Neurochemistry, University of Gothenburg^4^.

### Analysis of cognitive functions with mPACC3 scores

Three composite tests for measuring executive functions, along with assessments of two other cognitive domains—immediate learning and delayed recall—were used to compute the global composite scores for the three-test version of the Preclinical Alzheimer’s Cognitive Composite (PACC3). The PACC3 scores were derived using three distinct measures of executive functions—Animal Naming Test (PACC3-AN), Category Fluency Test (PACC3-CFL), and Trail-Making Test B (PACC3-TRLB)—and subsequently transformed into z-scores^5^. The modified Preclinical Alzheimer’s Cognitive Composite (mPACC3) scores were used for all associations pertaining to cognitive function. The mPACC was adapted to address sampling limitations during the COVID-19 pandemic with remote assessments^6^. The test battery included telemetric animal naming (mPACC3-AN), categoric fluency or letter fluency (mPACC3-CFL) tests. Additionally, we used in-person trail making test-B (mPACC3-TRL-B) results to include all available overlapping data without selection bias.

### CSF biomarkers

Cerebrospinal fluid (CSF) samples were collected via lumbar puncture in the morning after an 0-12 hour fast, as previously described^7^. CSF biomarkers were measured using the NeuroToolKit (NTK), a panel of exploratory robust prototype assays (Roche Diagnostics International Ltd, Rotkreuz, Switzerland). The following biomarkers were measured as markers of AD and related pathologies and quantified on the Cobas^®^ e 601 module (Roche Diagnostics International Ltd, Rotkreuz, Switzerland): Aβ42, pTau181, tTau, S100 calcium-binding protein B (S100B), and interleukin-6 (IL-6). The remaining biomarkers were assayed on the Cobas^®^ e 411 analyzer: Aβ40, neurofilament light protein (NFL), neurogranin, α-synuclein, glial fibrillary acidic protein (GFAP), chitinase-3-like protein 1 (YKL-40), and soluble triggering receptor expressed on myeloid cells 2 (sTREM2).

### FDG PET imaging

FDG PET imaging was performed using a Siemens EXACT HR+ tomograph in 3D mode at the Waisman Center Brain Imaging Laboratory, University of Wisconsin-Madison^8^. Prior to scanning, participants were screened to confirm glucose levels below 180 mg/dL. Imaging began with a 6-minute transmission scan for attenuation correction, followed by a 30-minute dynamic emission scan (6 × 5-minute frames) starting 30 minutes after administering a bolus injection of approximately 185 MBq of [18F]FDG. PET data were reconstructed using filtered back-projection, with corrections for random events, scatter, segmented attenuation, dead time, scanner normalization, and radioactive decay. The reconstructed images had a matrix size of 128 × 128 × 63 with voxel dimensions of 2.57 × 2.57 × 2.43 mm. PET images were processed using a custom MATLAB script. The reconstructed PET time series were smoothed with an isotropic 6 mm Gaussian kernel, realigned between frames using SPM12, and summed for the 30–60-minute post-injection period. Parametric standard uptake value ratio (SUVR) images were created by linearly aligning the summed PET image to the corresponding T1-weighted magnetic resonance imaging (MRI) and normalizing the intensities with cerebellar gray matter as the reference region. Regional mean FDG SUVR values were extracted from bilateral regions of interest (ROIs), including the frontal cortex, lateral temporal cortex (encompassing the middle and superior temporal gyri), supramarginal and angular gyri, precuneus, posterior cingulate, and hippocampus. The selection of these regions for analysis was based on earlier studies that identified reduced FDG uptake in individuals with AD dementia^9,10^.

### Shotgun sequencing

DNA library samples were prepared using Illumina NexteraXT library preparation kit. Quality and quantity of the finished libraries were assessed using an Agilent bioanalyzer and Qubit® dsDNA HS Assay Kit, respectively. Libraries were standardized to 2nM. Paired-end, 150 bp sequencing was performed using the Illumina NovaSeq6000, producing 2x150bp paired-end reads. Images were analyzed using the standard Illumina Pipeline (v1.8.2). Low-quality reads were filtered out from raw DNA reads using Trimmomatic (v0.39) with parameters (SLIDINGWINDOW:4:20 MINLEN:50). To identify and eliminate host sequences, reads were aligned against the Homo sapiens genome (GRCh38, Rel109) using bowtie2 (v2.3.4) with default settings, and microbial DNA reads that did not align with the human genome were identified using samtools (v1.3) (samtools view -b -f 4 -f 8). Shotgun metagenomic data are available from the Sequence Read Archive (SRA) under accession number PRJNA1271016.

### Proteomics and phosphoproteomics

Flash frozen brain cortices from 4 PS19 P301S male mice were homogenized in extraction buffer (25mM Tris HCl, 150mM NaCl, 10mM EDTA, 10mM EGTA, 1mM DTT, protease and phosphatase inhibitors) using ceramic beads and a bead-beater. Lysates were centrifuged at 16,000 x g for 20 min at 4°C and supernatants were collected. 200µL lysates were transferred to 2mL low-protein binding microcentrifuge tubes and 800µL of LC-MS grade methanol was added (80% final v/v) for BSL2 inactivation in biosafety cabinet by vortexing and incubation at room temperature for 5min. Subsequently 250µL chloroform was added, followed by heavy vortexing and 600µL water with heavy vortexing to facilitate protein precipitation and two-phase partitioning. Samples were spun down for 1min at max speed at room temperature and upper-phase was removed carefully not to disturb the protein interphase. 1mL methanol was added, samples were vortexed and spun for 4 minutes to pellet the protein, supernatant was removed and another round of protein extraction on second 200µL aliquot of lysate was done in the same tube to a total volume of 400µL lysate. Two additional cold acetone washes were performed by repeating aforementioned vortex and centrifuge steps in between and finally pellets were air dried to completion. Dried pellets were re-solubilized in 65µL of 8M Urea in 50mM NH4HCO3 (pH8.5) and 5µL was taken for BCA protein measurement. Subsequently 100µg of protein in 30µL 8M urea and 50mM NH4HCO3 (pH8.5) per each sample was taken for tryptic/LysC digestion where the samples were first diluted with 5μl of 25mM DTT and 25mM NH4HCO3 (pH8.5) to 120µL final volume for the reduction step, which was carried out for 15min at 56°C. After cooling on ice to room temperature 6μL of 55mM CAA (chloroacetamide) was added for alkylation where samples were incubated in darkness at room temperature for 15 minutes. This reaction was quenched by 16μL addition of 25mM DTT. Subsequently 40uL of trypsin/LysC solution consisting of 100ng/μL 1:1 Trypsin [Promega]:LysC [FujiFilm] mix in 25mM NH4HCO3 along with 18μL of 25mM NH4HCO3 (pH8.5) was added to the samples for a final 200µL volume. Digests were carried out overnight at 37°C then subsequently terminated by acidification with 30µL of 2.5% TFA (Trifluoroacetic Acid) to 0.32% final plus 3µL of 10% HFBA (Heptafluorobutyric acid) to 0.13% final and samples were desalted and concentrated using 10mg/1ml Strata™-X 33µm Polymeric Reverse Phase SPE cartridges [Phenomenex] per manufacturer protocol. Eluates in 70%:28%:2% acetonitrile:water:formic acid were dried to completion then reconstituted in 100µL of 100mM TEAB (Triethylammonium Bicarbonate) for TMT labelling with 40µL [20µg/µL] of reagent [TMT 10plex™ Labeling Reagent Set Lot#AA401548A from Thermo Scientific in 100% acetonitrile] done at room temperature for 1hr with intermittent gentle vortexing every 15min. The labeling reaction was terminated with 8µl addition of 5% hydroxylamine [0.27% final] and 15min incubation at room temperature. ∼1/37th (4µL, ∼2.6µg) of each individually labelled sample was taken and pooled together for a test pool run to evaluate complexity and mixing ratio. Post test-pool evaluation, master-pool sample was generated where each biological sample was present at the same total protein mass (∼1mg total protein in ∼1.1mL total volume), acidified to 1% formic acid (pH∼3), frozen (-80⁰C) then dried to almost completion in the speed vac. Subsequently resolubilized in 600µL of 5% acetonitrile, 0.3% TFA and 0.2% HFBA for solid-phase extraction using 30mg Strata™-X 33µm Polymeric Reverse Phase SPE cartridge [Phenomenex] per manufacturer protocol. 1/24th of eluate was used for the total protein profiling and the rest was retained for the phosphoenrichment where the eluate in 75/23/2% Acetonitrile/H2O/Formic acid was dried to completion in the speed-vac then brought back to 200µL total volume with 1M glycolic acid in 80% Acetonitrile and 5% Trifluoroacetic acid immediately before phosphoenrichment, which was carried out in two stages, first using 5mg of titanium dioxide magnetic microparticles [MagReSyn® TiO2 from ReSyn Biosciences] then unbound material was enriched with 2mg of Zirconium-ion (Zr4+) functional magnetic microparticles [MagReSyn® Zr-IMAC from ReSyn Biosciences] according to manufacturer protocol. Eluates from sequential enrichments were combined and dried in the speed-vac to completion. Sample handling for both enrichments were according to the manufacturer protocols. Enriched samples were solubilized in 35µL of 0.3% TFA, 0.2% HFBA and 3% ACN then desalted using 10µL Pierce® C18 Tips [Thermo Fisher Scientific] according to manufacturer protocol. Eluates in 75%:25%:0.1% acetonitrile:water:TFA acid (v/v) were dried to completion in the speed-vac and reconstituted in 15µL of 0.1% formic acid and 5% acetonitrile for injection on the instrument.

### NanoLC-MS/MS

Peptides were analyzed on Orbitrap Fusion™ Lumos™ Tribrid™ platform, where 3µL [∼2.4µg] total proteome or 3µL [1/5th] total enriched phospho-proteome was injected using Dionex UltiMate™3000 RSLCnano delivery system [ThermoFisher Scientific] equipped with an EASY-Spray™ electrospray source (held at constant 50°C). Chromatography of peptides prior to mass spectral analysis was accomplished using capillary emitter column [PepMap® C18, 2µM, 100Å, 500 x 0.075mm, Thermo Fisher Scientific]. NanoHPLC system delivered solvents A: 0.1% (v/v) formic acid , and B: 80% (v/v) acetonitrile, 0.1% (v/v) formic acid at 0.30 µL/min to load the peptides at 2% (v/v) B, followed by quick 2 minute gradient to 5% (v/v) B and gradual analytical gradient from 5% (v/v) B to 62.5% (v/v) B over 203 min when it concluded with rapid 10 minute ramp to 95% (v/v) B for a 9 minute flash-out. As peptides eluted from the HPLC-column/electrospray source survey MS scans were acquired in the Orbitrap with a resolution of 60,000, max inject time of 50ms and AGC target of 1,000,000 followed by HCD-type MS2 fragmentation into Orbitrap (36% collision energy and 30,000 resolution) with 0.7 m/z isolation window in the quadrupole under ddMSnScan 1 second cycle time mode with peptides detected in the MS1 scan from 400 to 1400 m/z with max inject time of 54ms and AGC target of 125,000; redundancy was limited by dynamic exclusion and MIPS filter mode ON.

### Protein enrichment

Protein enrichment was performed using differentially regulated hits (>±30% FC) that are statistically significant (FDR *P*<0.05) on Metascape^11^ to obtain direct overlaps, functional overlaps, and pathway overlaps between 5XFAD and PS19P301S models. To sort phosphoproteome by interacting kianses, we enriched top-25 (>30% FC) phosphoproteins using Kinase Enrichment Analysis 3 (KEA3)^12^ and cell compartment enrichment was performed using Enrichr^13–16^.

---

**Extended Data Figure 1.**
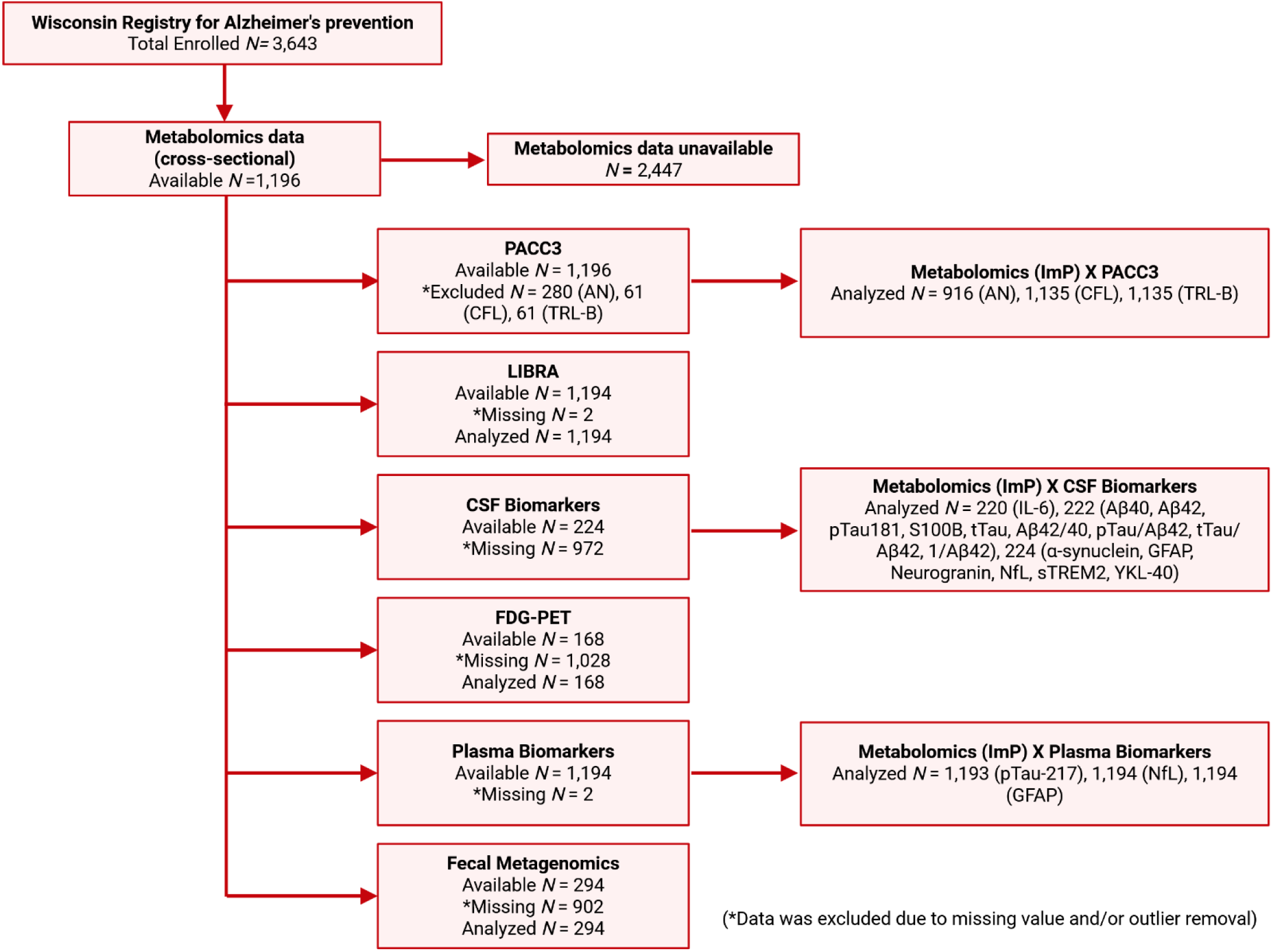
CONSORT flow diagram summarizing demographic information for cross-sectional assesment of associations between ImP and ADRD phenotypes.

**Extended Data Figure 2.**
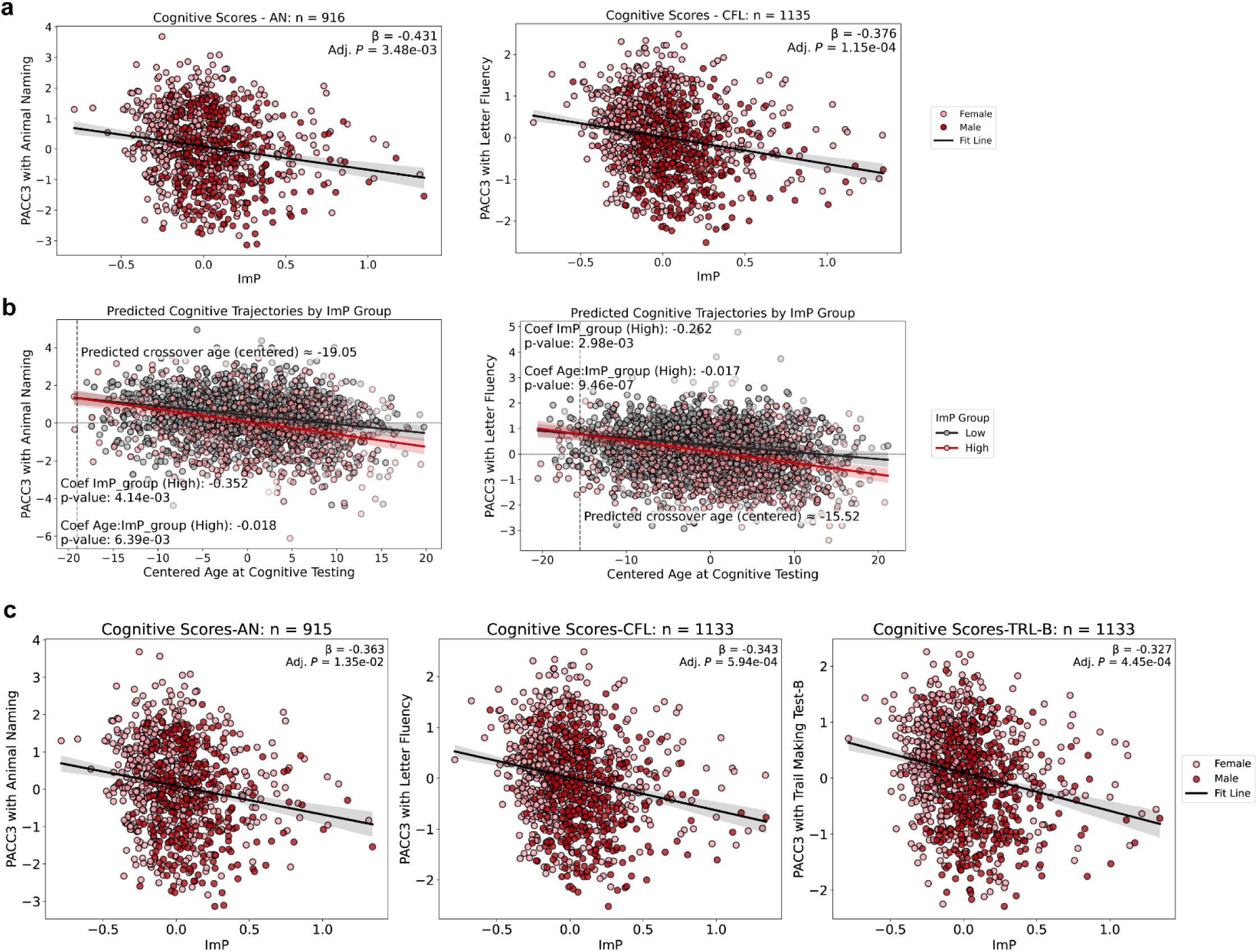
Plasma ImP is associated with lowered mPACC3 cognitive scores. a, Plasma ImP is negatively associated with mPACC3-AN and mPACC3-CFL. **b,** Modeled cognitive trajectories over age for mPACC3 scores (AN and CFL), with the vertical dashed line indicates predicted crossover age and the horizontal dashed line denotes zero on the scaled y-axis, comparing top (red) and bottom (gray) ImP quartiles. **c,** Plasma ImP is associated with pre-clinical cognitive scores agnostic to comorbid LIBRA scores as a covariate.

**Extended Data Figure 3.**
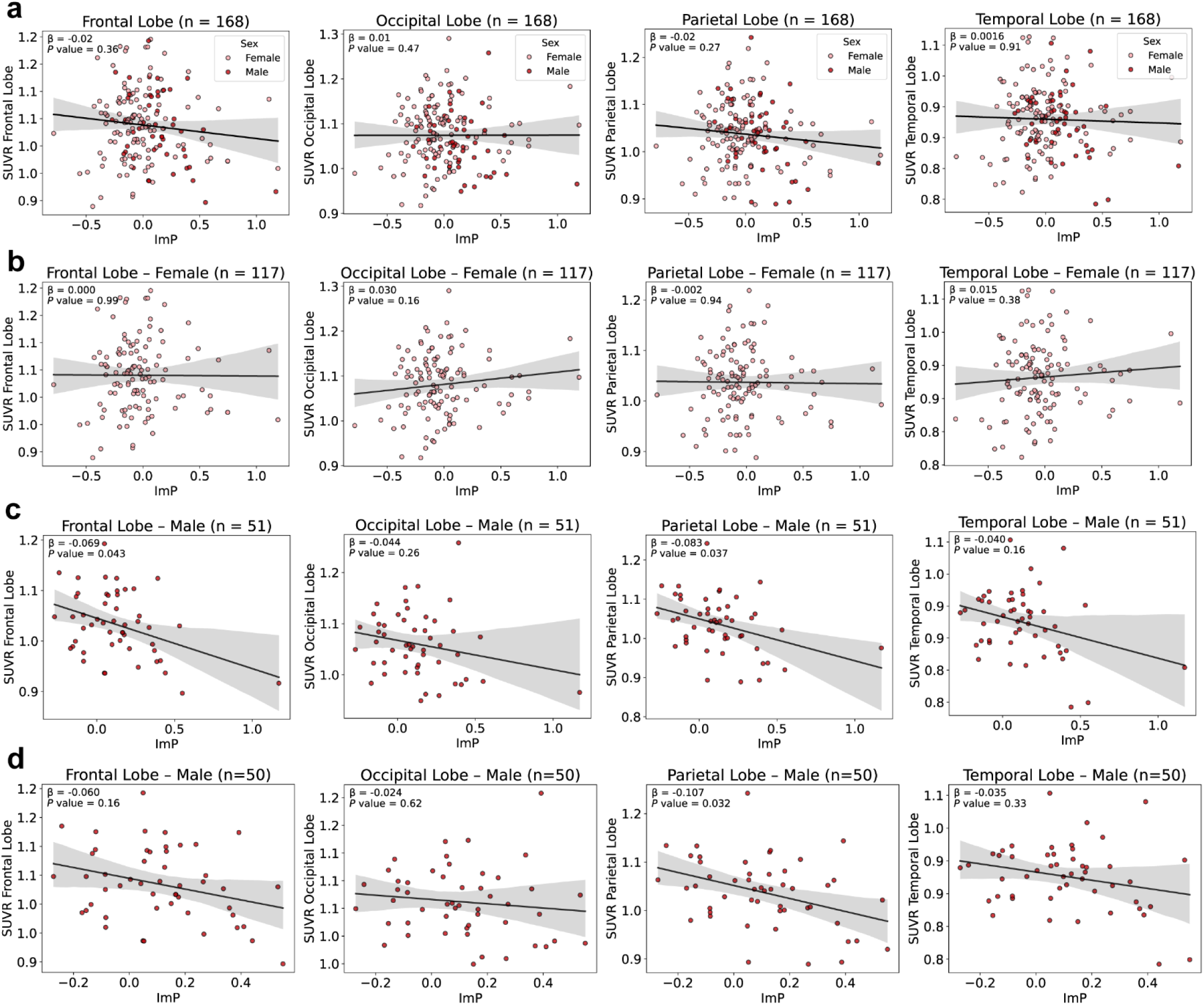
Plasma ImP levels are not significantly associated with FDG SUVR. a, Plasma ImP levels are not significantly associated with FDG SUVR in frontal, occipital, parietal and temporal lobes. **b,** Associations between FDG SUVR and ImP are not significant when stratified to female sex. **c,** FDG SUVR in males is significantly lowered in frontal and parietal lobes, but not in occipital and temporal lobes. **d,** FDG SUVR of males with outliers removed, shows statistically significant association only in parietal lobe.

**Extended Data Figure 4.**
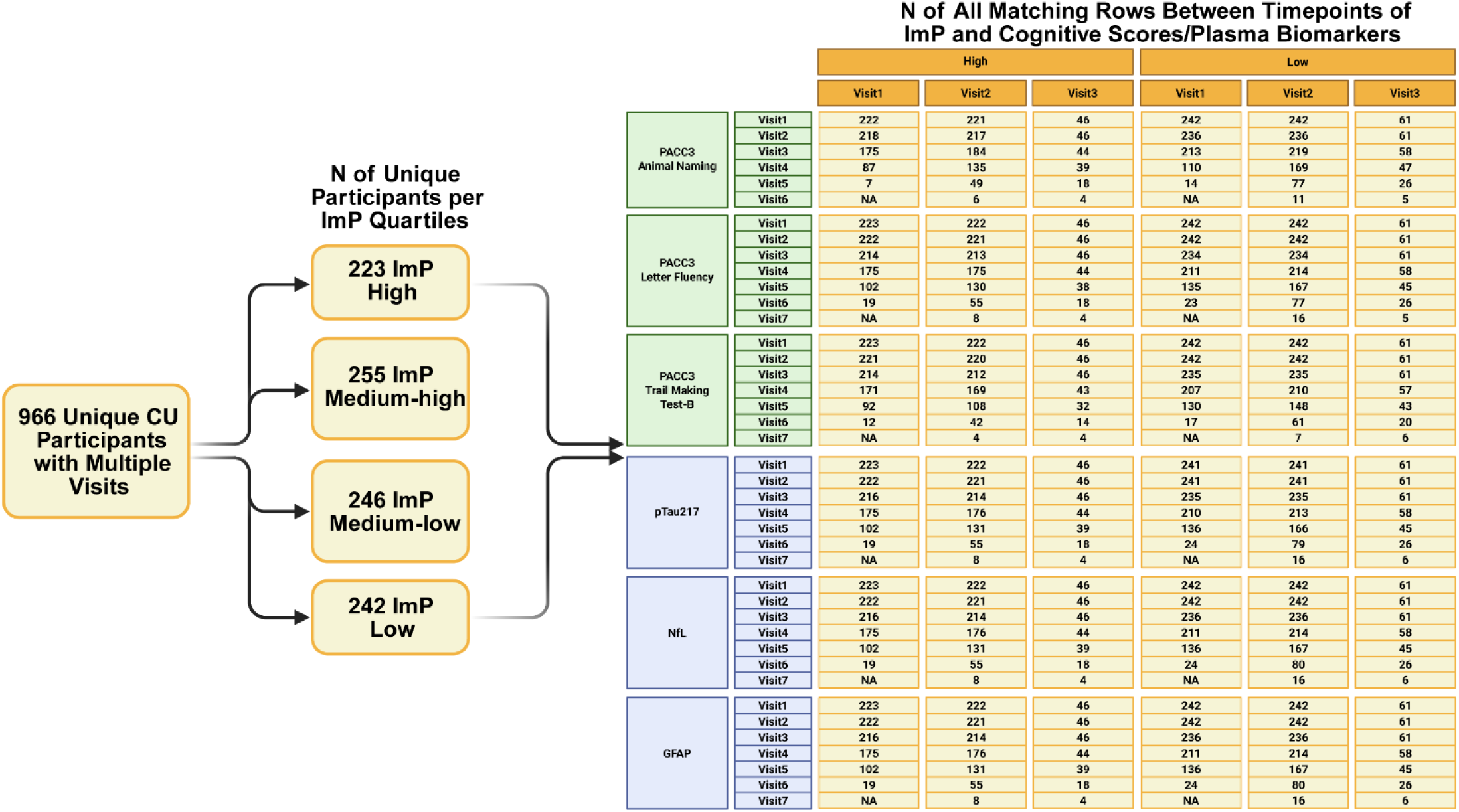
Demogrpahic table for stratification of WRAP ADRC cohort by quartiles based on plasma ImP levels for longitudinal analysis with distributions of number of subjects represented by outcome variables.

**Extended Data Figure 5.**
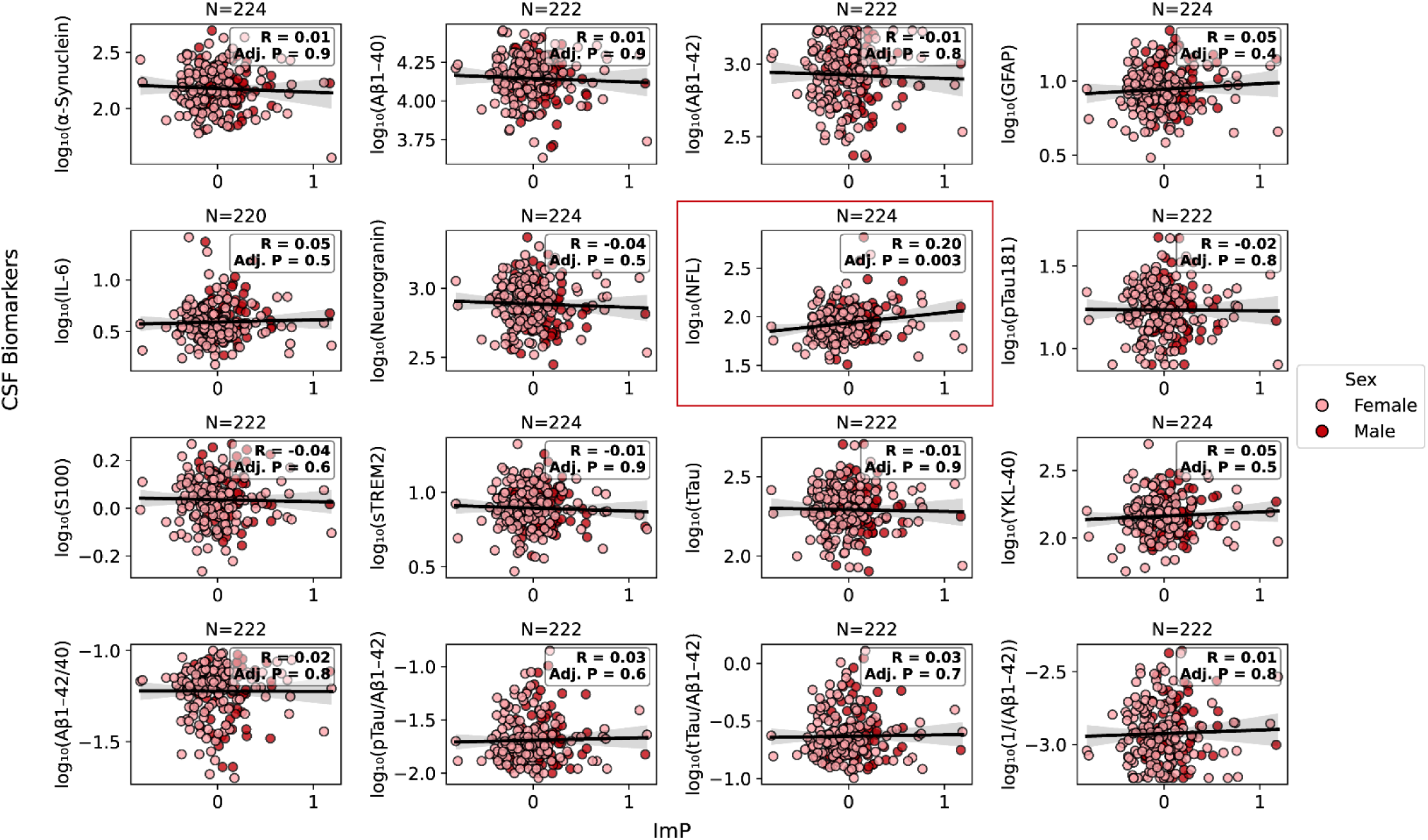
Plasma ImP levels are significantly associated with CSF neurofilament light chain levels (red box) but not other CSF biomarkers of ADRD.

**Extended Data Figure 6.**
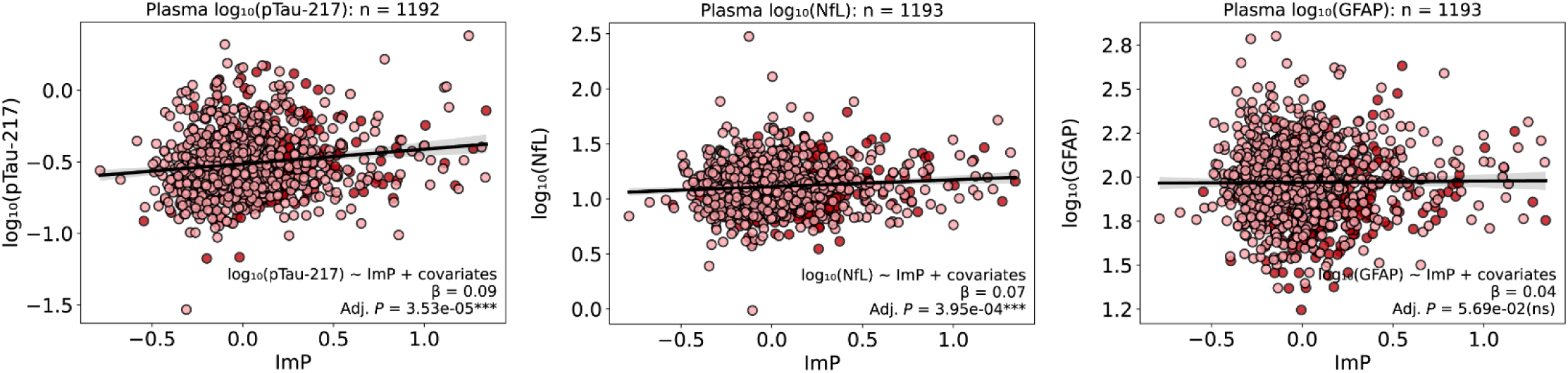
Plasma ImP is associated with plasma biomarkers agnostic to comorbid LIBRA scores as a covariate.

**Extended Data Figure 7.**
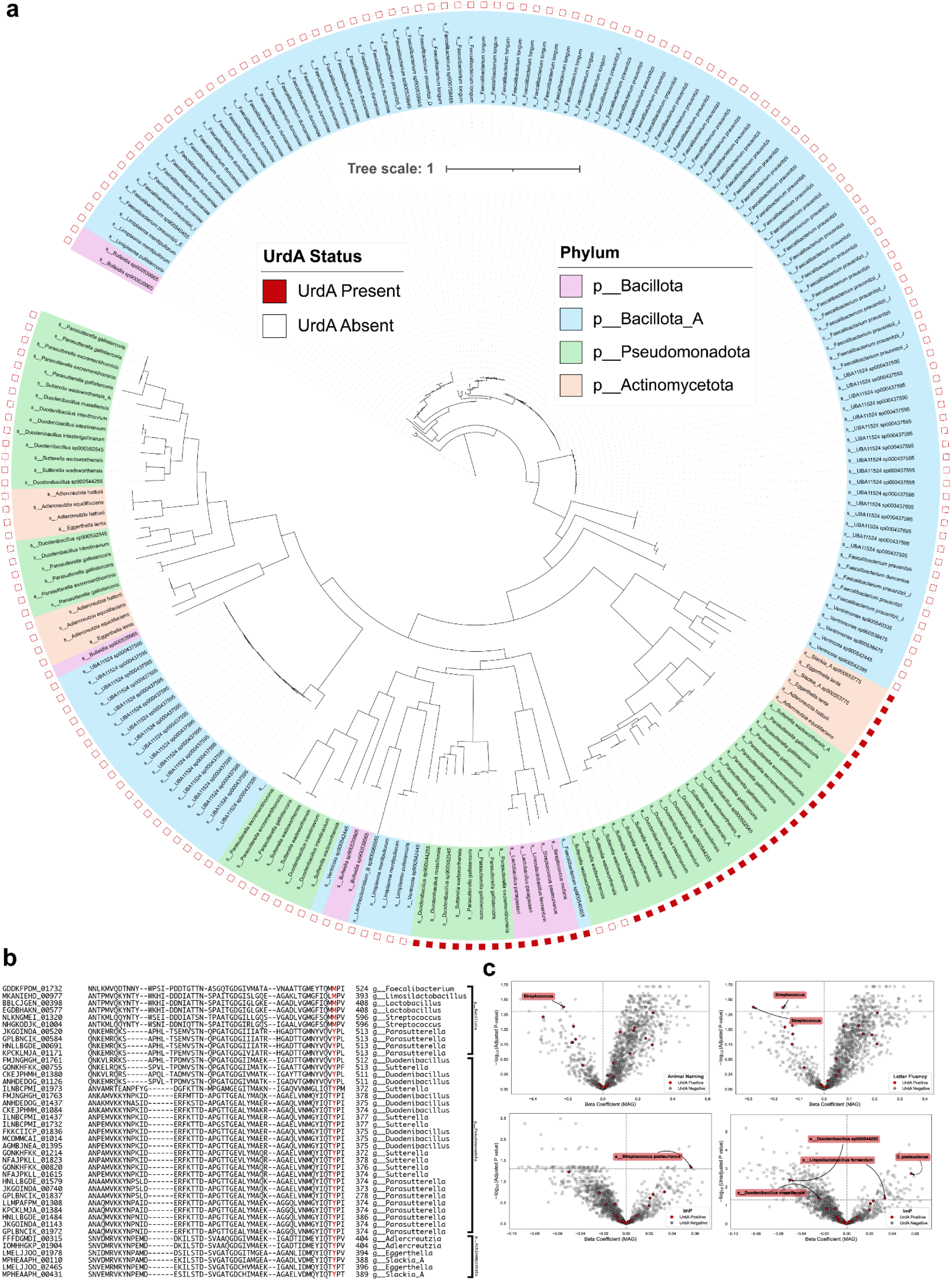
Fecal metagenomics revealed several gut bacterial taxa harboring putative *urdA* orthologs in *N*= 294 samples. a, Full sized phylogenetic tree depicting the various MAGs within MARS cohort (N = 294). Each terminal branch represents a species, colored by phylum. The outer layer of the tree indicates the presence (red) or absence (white) of the *urdA* gene. **b,** Representative *urdA* sequences marked with 373^rd^ amino acid in red. **c,** mPACC3 scores AN and CFL are negatively associated with *urdA^+^* taxa (Top). *Streptococcus pasteurianus* carrying putative *urdA* ortholog is positively associated with plasma ImP levels (Bottom).

**Extended Data Figure 8.**
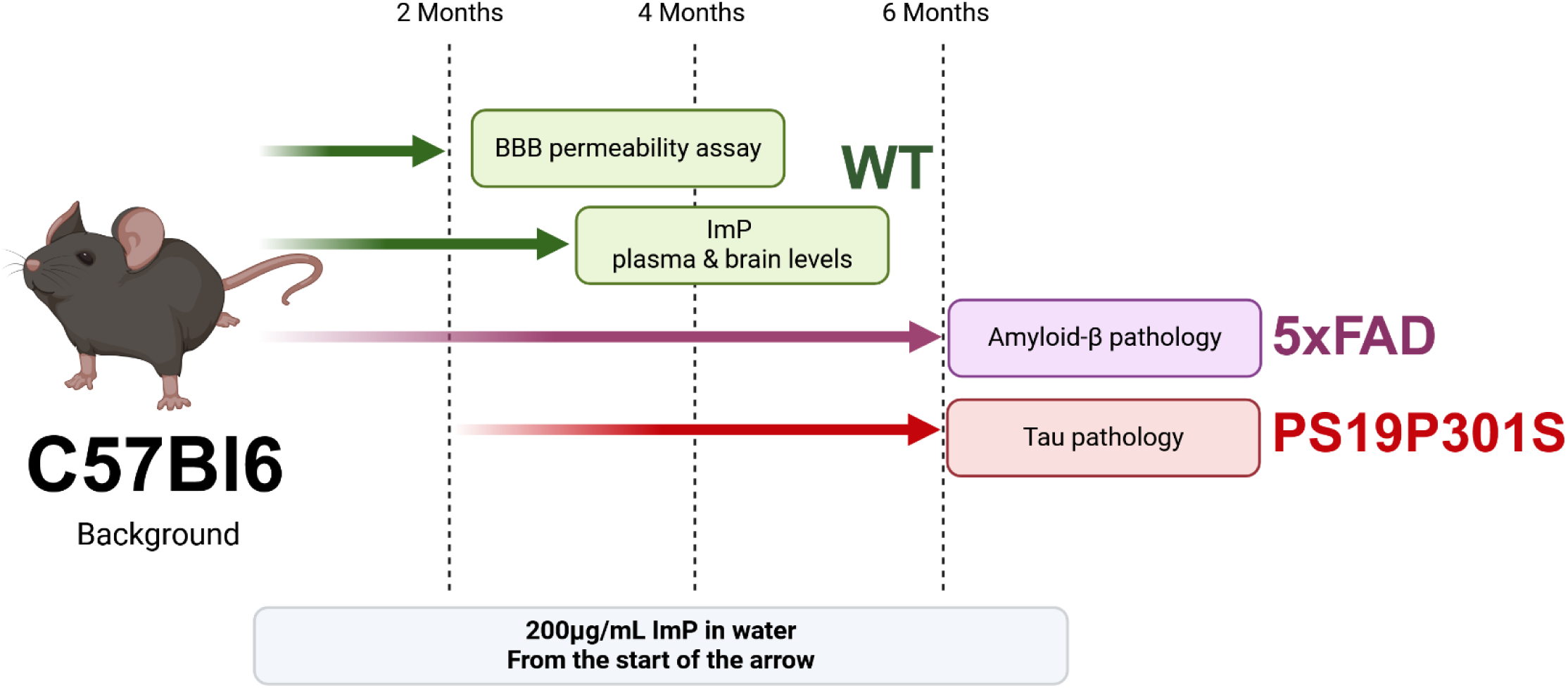
Mouse models of C57Bl6 background were employed to tests causal effects of ImP on ADRD-like pathology. Mice were treated with ImP orally through water from the starting point of arrows color coded by WT, or transgenic 5xFAD and PS19P301S models.

**Extended Data Figure 9.**
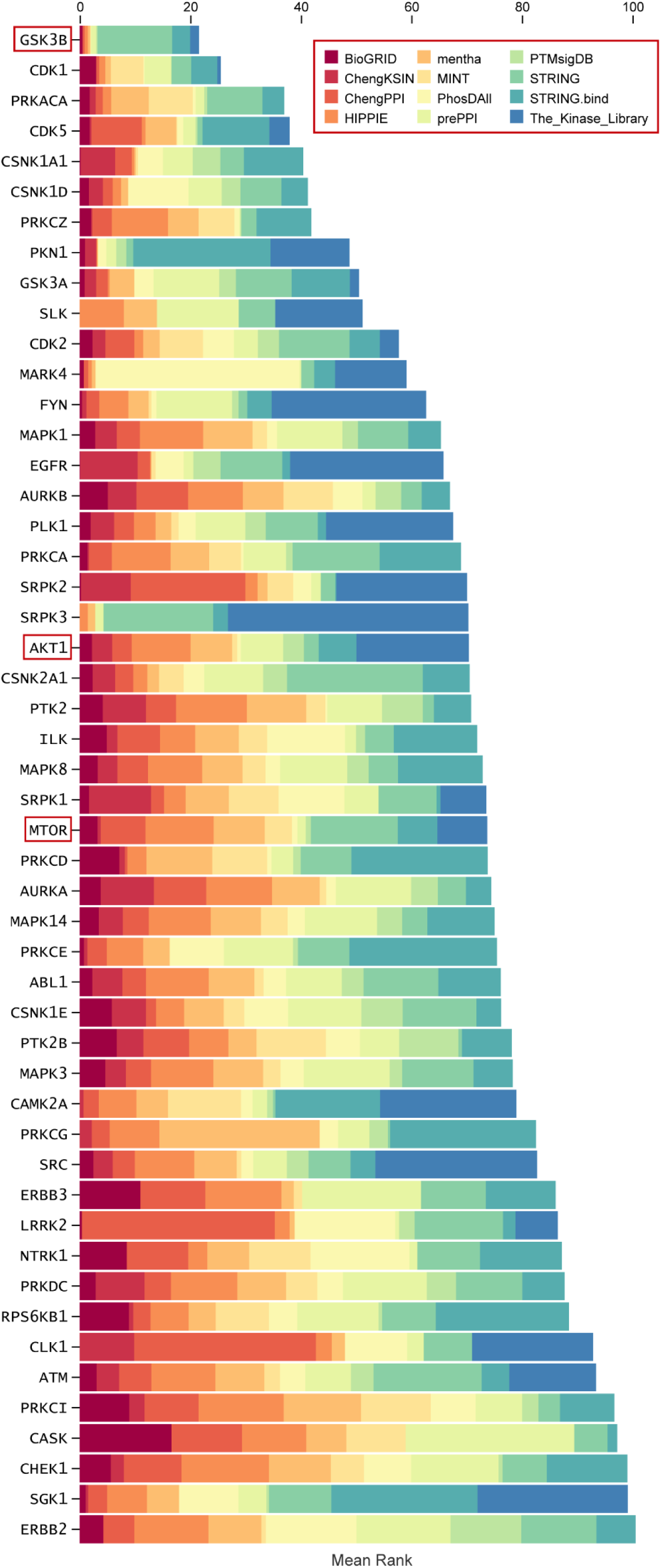
Kinase enrichment analysis of the top 25 differentially abundant phosphoproteins from PS19P301S cortical lysates, shown as stacked bar plots for top 100 highest-ranked kinases. Specific kinases of interest are marked with a red box.

**Extended Data Figure 10.**
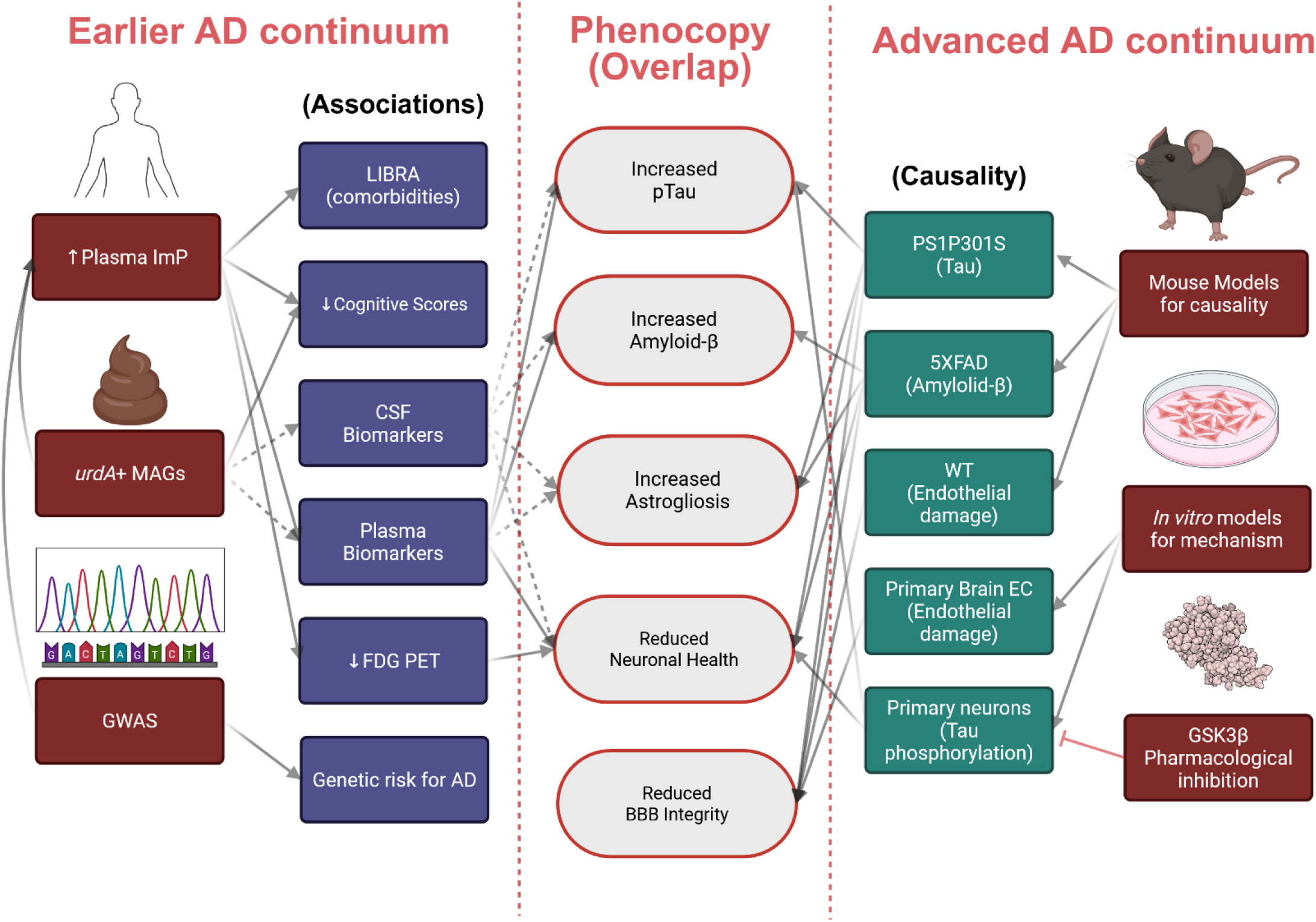
Simplified study schematic illustrating observations (dotted arrows indicate supporting trends) from the clinical datasets within earlier stages of AD continuum (cognitively unimpaired), except for GWAS study which associates ImP with AD. We identified that ImP is causal to exacerbated hallmark AD advanced-stage neuropathological features in controlled experiments featuring pre-clinical models. Overall, we identify multiple lines of evidence linking the gut-bacterial metabolite ImP to AD.

