## Supplementary figures and images for "Gut bacterial metabolite imidazole propionate potentiates Alzheimer’s disease pathology"

### Figure 3. Panel a

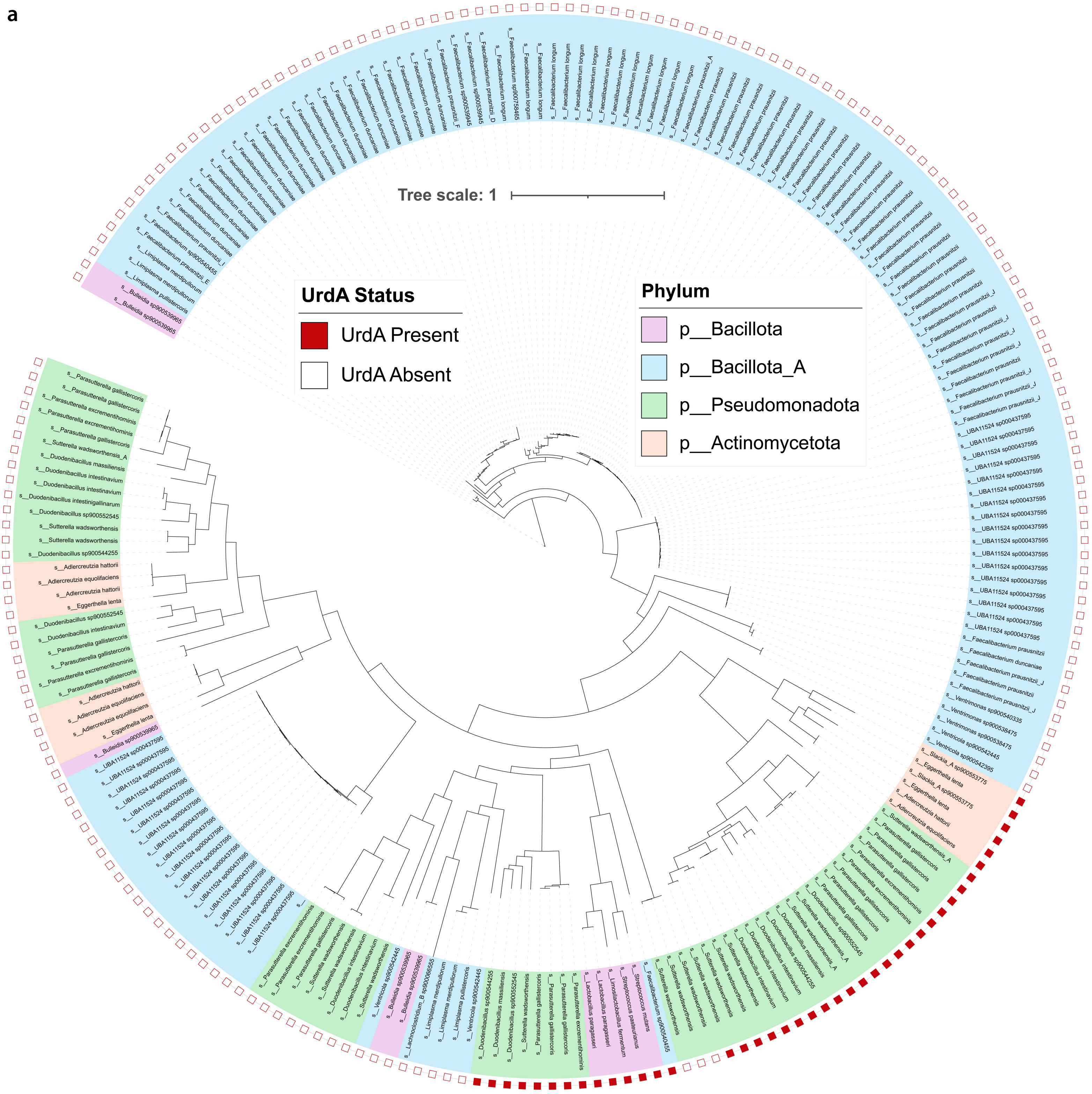
